# Delayed DNA replication in haploid human embryonic stem cells

**DOI:** 10.1101/2021.05.11.443666

**Authors:** Matthew M. Edwards, Michael V. Zuccaro, Ido Sagi, Qiliang Ding, Dan Vershkov, Nissim Benvenisty, Dieter Egli, Amnon Koren

**Affiliations:** Department of Molecular Biology and Genetics, Cornell University, Ithaca NY 14853, USA; Department of Pediatrics and Naomi Berrie Diabetes Center, Columbia University, New York, NY 10032, USA; Columbia University Stem Cell Initiative, New York, NY 10032, USA; The Azrieli Center for Stem Cells and Genetic Research, Department of Genetics, Silberman Institute of Life Sciences, The Hebrew University, Jerusalem 91904, Israel; Whitehead Institute for Biomedical Research, Cambridge, MA 02142, USA

## Abstract

Haploid human embryonic stem cells (ESCs) provide a powerful genetic system but diploidize at high rates. We hypothesized that diploidization results from aberrant DNA replication. To test this, we profiled DNA replication timing in isogenic haploid and diploid ESCs. The greatest difference was the earlier replication of the X chromosome in haploids, consistent with the lack of X chromosome inactivation. Surprisingly, we also identified 21 autosomal regions that had dramatically delayed replication in haploids, extending beyond the normal S phase and into G2/M. Haploid-delays comprised a unique set of quiescent genomic regions that are also under-replicated in polyploid placental cells. The same delays were observed in female ESCs with two active X chromosomes, suggesting that increased X chromosome dosage may cause delayed autosomal replication. We propose that incomplete replication at the onset of mitosis could prevent cell division and result in re-entry into the cell cycle and whole genome duplication.

**Highlights:**

- DNA replication timing of haploid ESCs profiled by WGS
- Extreme replication timing delays in haploid ESCs at unique genomic regions
- Replication delays associate with X-chromosome dosage in multiple systems
- Replication delayed regions correspond to underreplication in mouse polyploid cells

## Introduction

Embryonic stem cells are typically derived from *in vitro* fertilization of human oocytes. Alternatively, oocytes can be artificially activated to develop into blastocysts from which parthenogenetic stem cells, containing only maternally derived chromosomes, can be derived. Recently, fluorescence-activated cell sorting (FACS) has been used to isolate and maintain haploid parthenogenetic embryonic stem cells (h-pESCs) (Sagi et al., 2016). Haploid cells hold great promise as a tool for conducting loss-of-function genetic screens (Leeb et al., 2014; Yilmaz et al., 2020); for studying the stability of cell ploidy in development and disease, tolerance to ploidy changes, X chromosome inactivation and parental imprinting; and potentially for applications in regenerative and reproductive medicine (Li and Shuai, 2017; Yilmaz et al., 2016; Zhang et al., 2020). However, haploid cells are naturally unstable, experiencing high levels of spontaneous diploidization (Tarkowski et al., 1970; Yaguchi et al., 2018). Human h-pESCs diploidize at a rate of 3-9% every cell cycle (Sagi et al., 2016), which poses a major limitation for their use in genetic studies. This diploidization also raises fundamental questions regarding the stability of the haploid state in mammals.

Oocyte activation without fertilization can also occur *in vivo*, in which case it results in the development of ovarian teratomas (Linder et al., 1975; Stevens and Varnum, 1973) that consist mostly of diploid cells (Baker et al., 1998; Heskett et al., 2020; Stelzer et al., 2011), suggesting that diploidization occurs early in their development. Diploidization is not specific to parthenogenicity, as hydatidiform moles forming from the loss of the nucleus in a fertilized egg are found to be diploid as well (Fan et al., 2002). Similarly, most androgenetic stem cells derived from haploid eggs with only a paternal genome are also diploid (Sagi et al., 2019).

It has previously been shown that diploidization of mouse haploid stem cells occurs through two rounds of DNA replication without an intervening cell division – rather than by cell fusion (Leeb et al., 2012; Takahashi et al., 2014). While previous studies have focused on mitotic progression for explaining diploidization (Guo et al., 2017; Leng et al., 2017; Li et al., 2017), the fundamental reasons for the failure of haploid cells to normally progress through the cell cycle remain unknown. Haploid human stem cells show several notable differences compared to their diploid counterparts, including being smaller, having a larger surface area-to-volume ratio, and a higher proportion of mitochondrial relative to nuclear DNA. In addition, haploid cells have one active X chromosome (Xa), whereas diploid cells have one active and one inactive X chromosome (Xi). Therefore, relative to the autosomes, haploids have a ~two-fold higher expression of X-linked genes when compared to diploids (Sagi et al., 2016). Any of these differences, as well as the lack of homologous chromosomes *per se*, could underlie the instability of mammalian haploid cells.

Diploidization resembles polyploidization, which also involves whole genome duplication. Polyploid cells normally arise in the human liver, bone marrow and placenta, as well as across tissues of *Drosophila melanogaster*, in many plants, and in other organisms (Sagi and Benvenisty, 2017; Schoenfelder and Fox, 2015). Polyploidy is also common in cancer (Bielski et al., 2018). Interestingly, polyploidization is often accompanied by genomic regions of reduced relative DNA copy number. In *Drosophila* polyploid cells, underreplication is due to active inhibition of replication fork progression in a subset of late-replicating genomic regions. At least two negative regulators of DNA replication, SUUR (Suppressor of Under-Replication) and Rif1, have been implicated in this process (Makunin et al., 2002; Munden et al., 2018; Nordman et al., 2011; Nordman et al., 2014). More recently, the highly polyploid trophoblast giant cells (TGC) of the mouse placenta have also been shown to harbor unique regions of reduced copy number (Hannibal et al., 2014).

## Results

### Haploid embryonic stem cells show replication timing profiles characteristic of pluripotent stem cells

We hypothesized that defects in DNA replication could be the fundamental cause of haploid cell diploidization, for instance by incomplete DNA replication carrying over to mitotic failure. To test for differences in DNA replication timing between haploid and diploid cells, we generated genome-wide DNA replication timing profiles for isogenic parthenogenetic haploid (h) and diploid (d) cultures of two cell lines, pES10 and pES12 (Sagi et al., 2016) along with long-term stably diploid pES10 and pES12 (unsorted for ploidy; see **Methods**) and two control ESC lines derived by *in vitro* fertilization (CU-ES4 and CU-ES5). DNA replication results in differential DNA copy number across the genome, with earlier-replicating loci showing an increased DNA content. These fluctuations in copy number can be detected from whole genome sequencing of cell populations and used to generate high-resolution profiles of DNA replication timing (Ding et al., 2020; Koren et al., 2014). Accordingly, we sequenced genomic DNA and calculated DNA copy number (sequencing read depth) in 2.5Kb windows of uniquely alignable sequence, normalized by local GC content (Koren et al., 2014). We filtered-out copy number variants (CNVs) and outliers, then smoothed the data to generate DNA replication timing profiles (**Methods**).

To assess the quality of the data, we first analyzed the autocorrelation of the unsmoothed replication timing profiles, which is a measure of profile continuity. Long-range autocorrelation was observed in haploid, isogenic diploidized (hereafter referred to as “diploid” for simplicity), and control cells across all chromosomes (**Fig. S1A**). We then compared the replication timing profiles of the haploid, diploid and control cell lines, as well as 57 separately sequenced ESCs with indications of normal X chromosome inactivation (karyotypically XX and henceforth referred to as female) (Ding et al., 2020)). The correlations within and between haploid and diploid cell lines ranged from r=0.92 to r=0.96, while the correlations to control ESCs ranged from between r=0.77 to r=0.88 when compared to the separately-sequenced ESCs, r=0.81 to r=0.93 compared to the concurrently sequenced control ESCs, and r=0.94 to r=0.95 compared to the isogenic unsorted diploid ESCs (**Fig. S1B**). Thus, the replication profiles were highly reproducible and consistent with controls and previous measurements, with expected minor differences likely due to experimental protocols and genetic background. In particular, control cell lines are biparental and may have specific differences from parthenogenetic cell lines at imprinted regions; our analyses below focus on comparisons between the isogenic parthenogenetic haploid and diploid cell lines. Finally, visual inspection of the replication timing profiles showed a high degree of similarity in the profiles of haploid, diploid, and control ESCs along the vast majority of the genome (**Fig. 1A**), with some notable differences explored further below.

**Figure 1.**
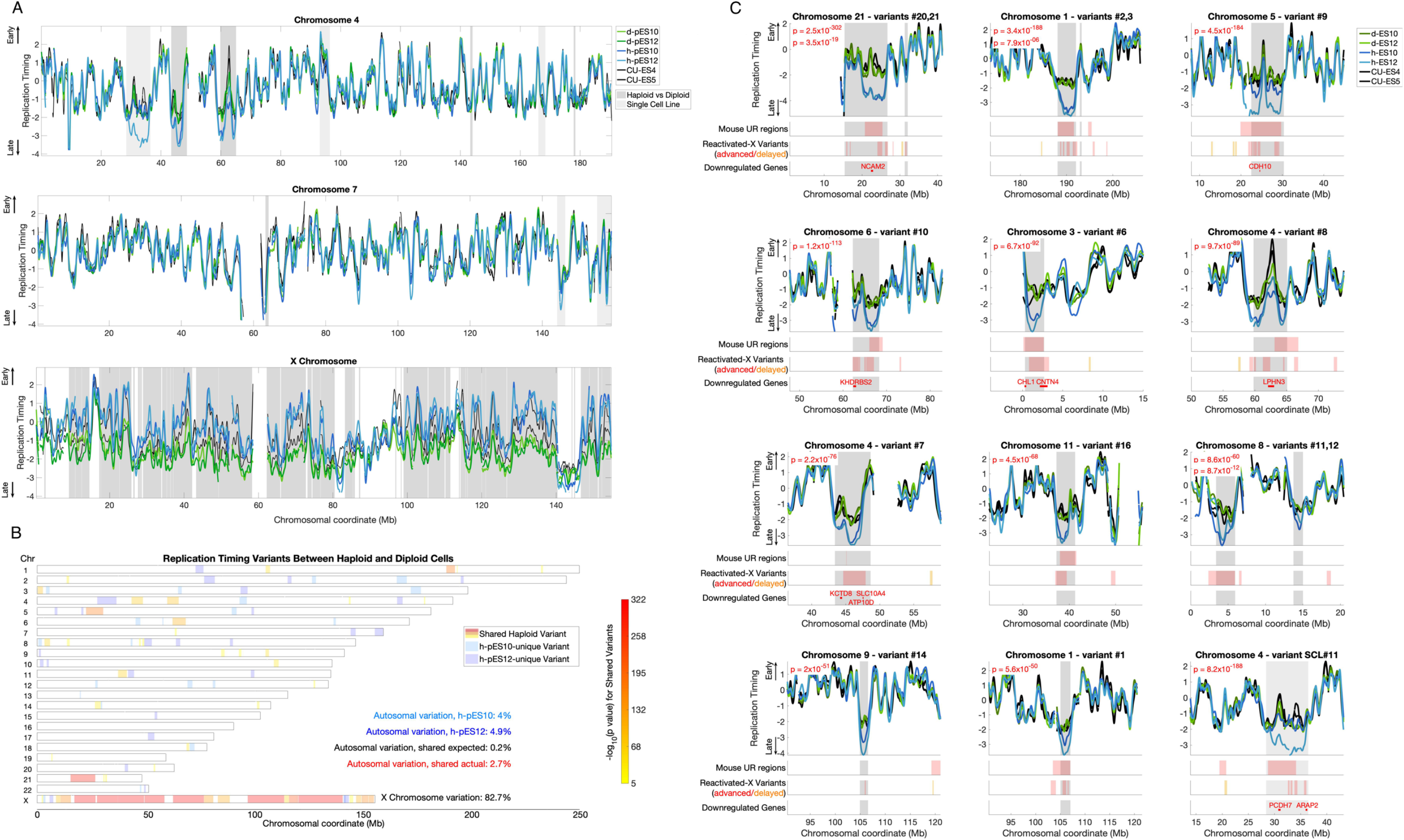
Significant replication timing variation between haploid and diploid ESCs. **A.** Representative replication timing profiles for haploid and diploid pES10 and pES12 and control ESCs. Profiles are z-score-normalized to a genome-wide average of zero and a standard deviation of one, such that positive and negative values indicate replication timing that is earlier and later than average, respectively, and the units corresponds to standard deviations (SD). Grey: regions of variation between haploid and diploid cells (light: one cell line; dark: shared in both cell lines). **B.** A genome-wide view of all replication timing variants. Shared variants are color-coded by p-value. The degree of shared haploid variation is significantly greater than expected given the extent of single cell line variation (chi-squared test p ≪ 10^−16^). **C.** The eleven most significant haploid-delayed variants, numbered by genomic location and ordered by p-value. Variants #21, #3, and #12 are each proximal to another, larger variant (represented by the second-listed p value in each panel). Variant SCL#11 (single cell line variant #11) is only delayed in a h-pES12, but nonetheless shows features common to other haploid-delayed variants (see text). Shown below each variant are the locations of all mouse UR regions (those found in all stages of development) and reactivated-X variants in the plotted interval of each panel. Reactivated-X delays are shown in red, while advanced regions are in orange. Downregulated genes are shown only within the replication timing variant borders (grey shades). Figure S2 shows all remaining variants.

### Significant replication timing variation between isogenic haploid and diploid human embryonic stem cells

Notwithstanding the high overall similarity in replication timing between haploid and diploid cells, we also observed sites of replication timing variation between the two cell ploidies or in individual haploid cell lines (**Fig. 1A**). To systematically identify and categorize such replication timing variants, we utilized one-way ANOVA tests on consecutive regions across the genome to identify consistent variation between the two haploid and the two diploid cell lines, and Student’s t-tests to identify regions in which only one haploid cell line was different than diploid cells (**Methods**). While differences specific to a single cell line could arise from genetic differences (Koren et al., 2014), somatic copy number alterations (see further below), or measurement noise, shared differences specifically reflect associations between cell ploidy and DNA replication timing.

To evaluate the specificity of the ANOVA test, we compared the extent of identified haploid-diploid replication timing variation across the autosomes to that of a sample permutation that disrupted the ploidy and genotype relationships (h-pES10 and d-pES12 compared to h-pES12 and d-pES10; **Methods**), revealing a 14.2-fold greater haploid-diploid variation than expected by default. The most extensive variation encompassed 82.7% of the X-chromosome across 19 distinct regions (86.8% when excluding PAR1 and evolutionary strata 4 and 5). While replication timing was highly correlated between haploid and diploid cells across the autosomes (r=0.92 to r=0.94), their correlations on the X chromosome were substantially lower (r=0.73 to r=0.76) (**Fig. S1B**). Autocorrelation of the X chromosome was also significantly higher in haploids compared to diploids (**Fig. S1A**). The mean replication timing of the X chromosome was much earlier in haploids (−0.29) compared to diploids (−1.6) (**Fig. 1A**). These results can be readily explained by X-chromosome inactivation in diploid cells: diploid pES10 and pES12 have an inactivate X chromosome (Sagi et al., 2016), which was shown before to replicate much later than the active X chromosome and without a well-defined replication timing program (Koren and McCarroll, 2014); this pattern of X chromosome replication is also presently observed in human ESCs (see below; **Fig. S3A,C**). Haploid cells, in contrast, only carry a single, active X chromosome. These results reaffirm the quality of our replication timing measurements and indicate that X chromosome inactivation, which leads to the largest gene expression differences between haploids and diploids (Sagi et al., 2016), is also reflected in DNA replication timing differences in haploids compared to diploids.

In addition to the large-scale variation on the X-chromosome, we also found 31 replication timing variants on the autosomes. These variants ranged from 392Kb to 11.3Mb in length (mean 2.3Mb), cumulatively covered 2.7% of the autosomes, and ranged in significance from p = 7.9 × 10^−6^ to p = 2.5 × 10^−302^ (**Fig. 1B,C, Fig. S1C**). We also identified 19 variants unique to h-pES10 and 24 variants that were specific to h-pES12. These cell line-specific variants provided a useful way to further evaluate the significance of having found as much shared haploid-diploid variation as we observed. Specifically, h-pES10 was variant from diploid cells across a total of 4.0% of the genome (1.3% specific to h-pES10 and 2.7% shared in both haploid cell lines) while h-pES12 was variant across 4.9% of the genome (2.2% cell-line specific).

Therefore, coincidental shared variation is expected to encompass 0.2% of the genome by chance (4.9% × 4%). However, we observed 2.7% of the genome to harbor shared replication timing variation between the two cell lines, a significant, 13.5-fold enrichment compared to expectation (chi-squared p ≪ 10^−16^; **Fig. 1B**). In addition, single cell line variants were less significant on average and showed much less correlation with other biological properties (see below). A notable exception is a large variant in h-pES12 on chromosome 4 (**Fig. 1C,** variant SCL#11), which was larger (by 2.55-fold) and much more significant than all other single cell line variants (p = 2.7 × 10^−149^, compared to the next strongest p-value of 2 × 10^−42^). Notwithstanding this exceptional variant, the strong enrichment of variation shared by both haploid cell lines suggests that the haploid state *per se*, and not sporadic variation between cell lines, is driving most of the replication timing variation that we identified. We conclude that human haploid ESCs have significant replication timing differences from their diploid counterparts on both the X chromosome and the autosomes. These results were reproduced in two additional replicate experiments (see further below).

### The replication of a subset of normally late-replicating genomic regions is further delayed in haploid cells

Of the 31 shared autosomal replication timing variant regions, 21 replicated later in haploid cells compared to diploids (“haploid-delayed”), while only 10 replicated earlier in haploids (“haploid-advanced”) (**Fig. 2A**). The haploid-delayed variants were larger than haploid-advanced variants (median size 2.01Mb compared to 836Kb, rank-sum p = 0.019), showed a greater replication timing difference compared to diploid cells (median of 1.58 SD from the mean, compared to 0.80, rank-sum p = 4.2 × 10^−4^), and were more significant (median p = 5.57 × 10^−50^ compared to p = 8.87 × 10^−10^, **Fig. 2B**). We ruled out the possibility that sites of replication delay could have a reduced copy number because of clonal or sub-clonal deletions (**Methods**; **Fig. S2A**).

**Figure 2.**
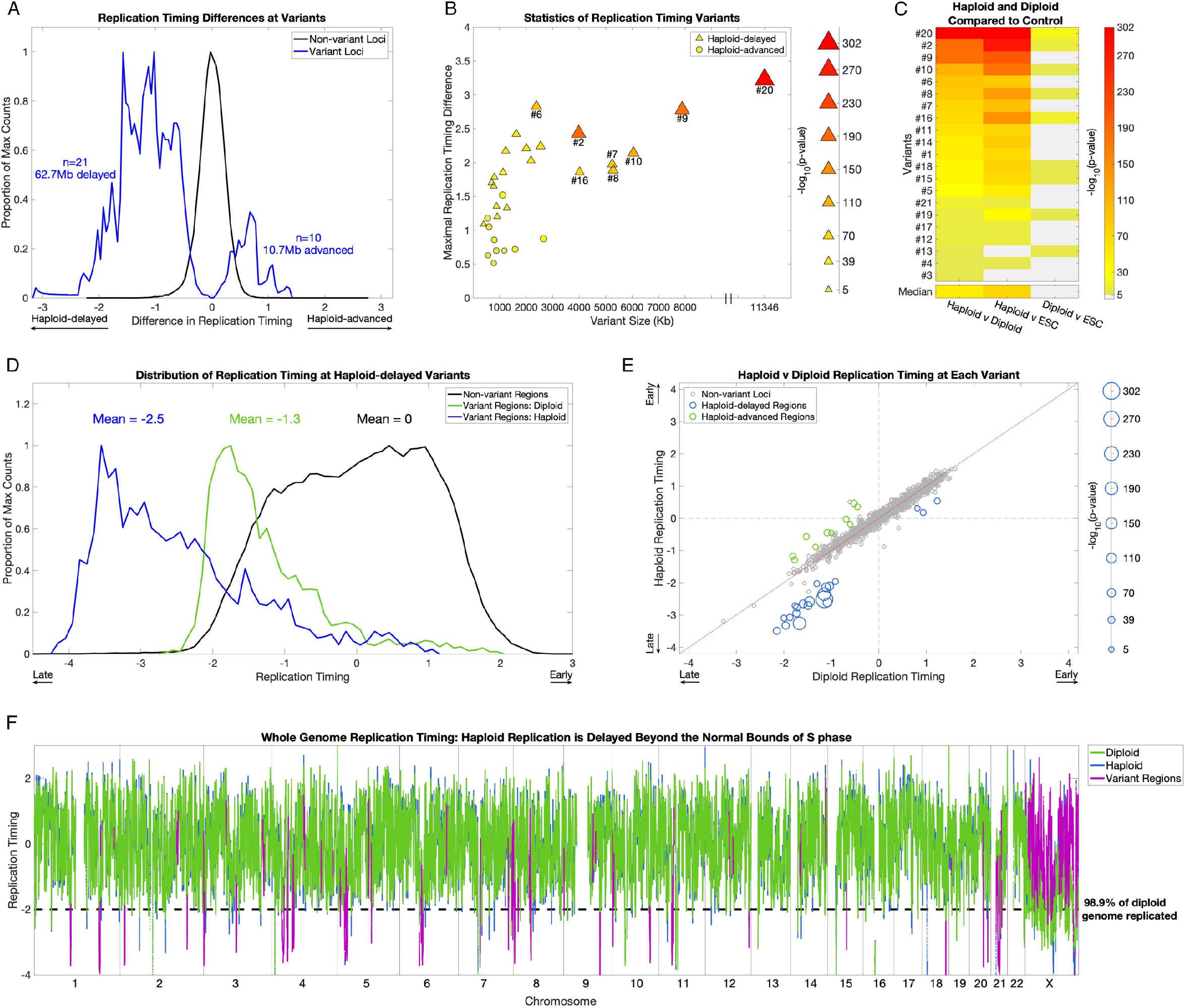
Replication timing in haploid cells is delayed beyond the normal S phase. **A.** Haploid-delays are the predominant form of replication timing variation. Distributions of the replication timing differences between haploid and diploid cells (mean of each pair of cell lines) in non-variant and variant regions. The distributions represent individual windows (2.5Kb) in each category. **B.** The size (scaled by the −log_10_(p-value)), absolute replication timing difference, and variation p-value of all 31 haploid-diploid replication timing variants. Haploid-delayed variants segregate from haploid-advanced variants on all three metrics, shown by the clustering of haploid-advanced (circles) in the bottom left, as well as their weaker significance. Conversely, haploid-delayed variants (triangles) are longer, have a greater replication timing difference from diploids, and are more significant. **C.** Replication timing variation is attributed to alterations in haploid cells. One-way ANOVA p-values between haploid, diploid, and control ESC lines (CU-ES4 and CU-ES5, shown as “ESC”) at haploid-delayed variants, sorted by p-value of haploid vs diploid comparisons (leftmost column). Light gray: not significant. Nineteen of the 21 haploid-delayed variants are also significantly different from controls (middle column), in contrast to only nine variants that vary between diploid cells and ESCs (rightmost column). In all but one case (variant #13), haploid-ESC variation is more significant than diploid-ESC variation, thus haploids, not diploids, have the exceptional replication timing values. **D.** Replication timing variants are normally late-replicating regions that are further and severly delayed in haploids. Distributions of replication timing for diploid cells in non-variant regions (i.e. normal replication timing distribution; black), and for diploid (green) and haploid (blue) cells in haploid-delayed variant regions. The distributions represent individual windows (2.5Kb) in each category. Haploid-delayed variants replicate late (on average, 1.3 SD later than the genome mean) in diploid cells, and considerably later (−2.5 SD later than the mean and beyond the bounds of normal S phase) in haploid cells. **E.** The majority of haploid-delayed variants are already late-replicating in diploid cells. Scatter plot of haploid vs diploid replication timing across the genome. Non-variant loci were defined by first removing variants, then binning the genome into 2.3Mb regions (mean size of all replication timing variants). Haploid-delayed variants (blue) replicate late in diploid cells and even later in haploids. The size of data points is scaled by −log_10_(p-value). The three dots (corresponding to variants #3, #4, and #13) with both early haploid and diploid replication had the poorest p-values of all haploid-delayed variants (p = 7.9×10^−6^, 2.3×10^−10^, and 2.8×10^−10^). **F.** Replication in haploid cells is delayed beyond the bounds of S phase. The dashed line indicates a replication timing of 2 SDs below the mean, when 98.9% of the genome has already completed replication. Eighteen of the 21 replication timing delays (all but the three least significant variants #3, #4, and #13) extended beyond this value.

To resolve whether variation was attributed to replication timing changes in the haploid cells, the diploid cells, or both, we compared the replication timing profiles to control ESCs. The haploid cell lines showed significant differences from control ESCs at 19 out of 21 haploid-delayed variants (p-values ranging from 1.9 × 10^−6^ to 2.5 × 10^−302^, **Fig. 2C**), compared to just nine variants for the diploid cell lines (p-values ranging from 1.5 × 10^−6^ to 5.7 × 10^−29^), which in every case were substantially weaker than the corresponding variation between haploid and diploid cells (**Fig. 2C**). The greater difference from controls of haploid compared to diploid cells suggests that replication timing delays are largely due to haploid-specific replication timing changes (justifying their designation as “haploid-delayed”). This comparison had a less clear interpretation at haploid-advanced variants, where we found a mixture of effects with greater ambiguity and more subtle differences between haploid and diploid cells (**Fig. S2B,C**). Taken together, these results indicate that delayed replication in haploid cells is the predominant replication timing difference between haploid and diploid cells.

Importantly, replication timing at haploid-delayed variants was already late in diploid cells, with a mean replication timing of 1.2 SDs later than the autosome-wide mean. In haploid cells, replication was further delayed to an average of 2.5 SDs, and a maximum of 4.1 SDs, later than the mean (**Fig. 2D**). This brings the average replication timing of these variant regions in haploids cells to later than 99.8% of all other autosomal loci (**Fig. 2D-F**). Given that 98.6% of the diploid genome replicates by 2 SDs later than the mean, a delay of 4.1 SDs extends S-phase by an estimated 52.5%. In ESCs, the duration of S-phase is approximately 8 hours, while G2 spans ~4 hours (Becker et al., 2006); an extension of 52.5% translates into a delay of 4.2 hours, potentially extending replication into late G2 phase and even into mitosis. Only three haploid-delayed variants (#3, #4, and #13) replicated earlier than the autosomal mean in haploid (as well as diploid) cells, and these were the three least significant variants (**Fig. 1, Fig. 2E, Fig. S1C**). Thus, replication timing variation between haploid and diploid ESCs comprises predominantly delays in already late-replicating regions, rendering these regions extremely late replicating and greatly extending the bounds of S phase, possibly into mitosis.

### Replication delays in haploid cells occur in quiescent, unorganized heterochromatin

Given that haploid-delayed variants are late-replicating and show severly delayed replication, we considered whether they correspond to fragile sites – genomic regions that are prone to double stranded breaks and are often late-replicating. However, neither haploid-delayed nor haploid-advanced variants significantly overlapped various classes of fragile sites (**Methods**; **Fig. S2D**). Haploid-delayed variants had a lower gene density than expected, even after accounting for the general sparsity of genes in late-replicating genomic regions (**Fig. 3A**, p = 0.011). The 181 genes within haploid-delayed variants were enriched for the gene ontology terms keratinization (23 genes, FDR = 1.49 × 10^−19^; **Table S1**), which was entirely attributed to the KRTAP gene cluster in variant #21, and cell-cell adhesion via plasma-membrane adhesion molecules (nine genes across six variants, FDR = 3.34 × 10^−3^). There was a significant enrichment (p = 0.03, **Fig. 1C, Fig. 3A**) within haploid-delayed variants for genes previously found to be downregulated in haploid cells (Sagi et al., 2016), as well as a strong correlation (r = 0.72) between the magnitude of haploid replication delay and the extent of gene expression downregulation (**Fig. 3B**). Downregulated genes within haploid-delayed variants were modestly enriched for cell-cell adhesion genes (FDR = 0.014) and components of the membrane (FDR = 0.033; **Table S1**). Similarly, haploid-advanced regions were enriched for genes upregulated in haploid cells (**Fig. S1D**; 11 genes compared to an expected 0.48; p = 3.7 × 10 × 10^−31^).

**Figure 3.**
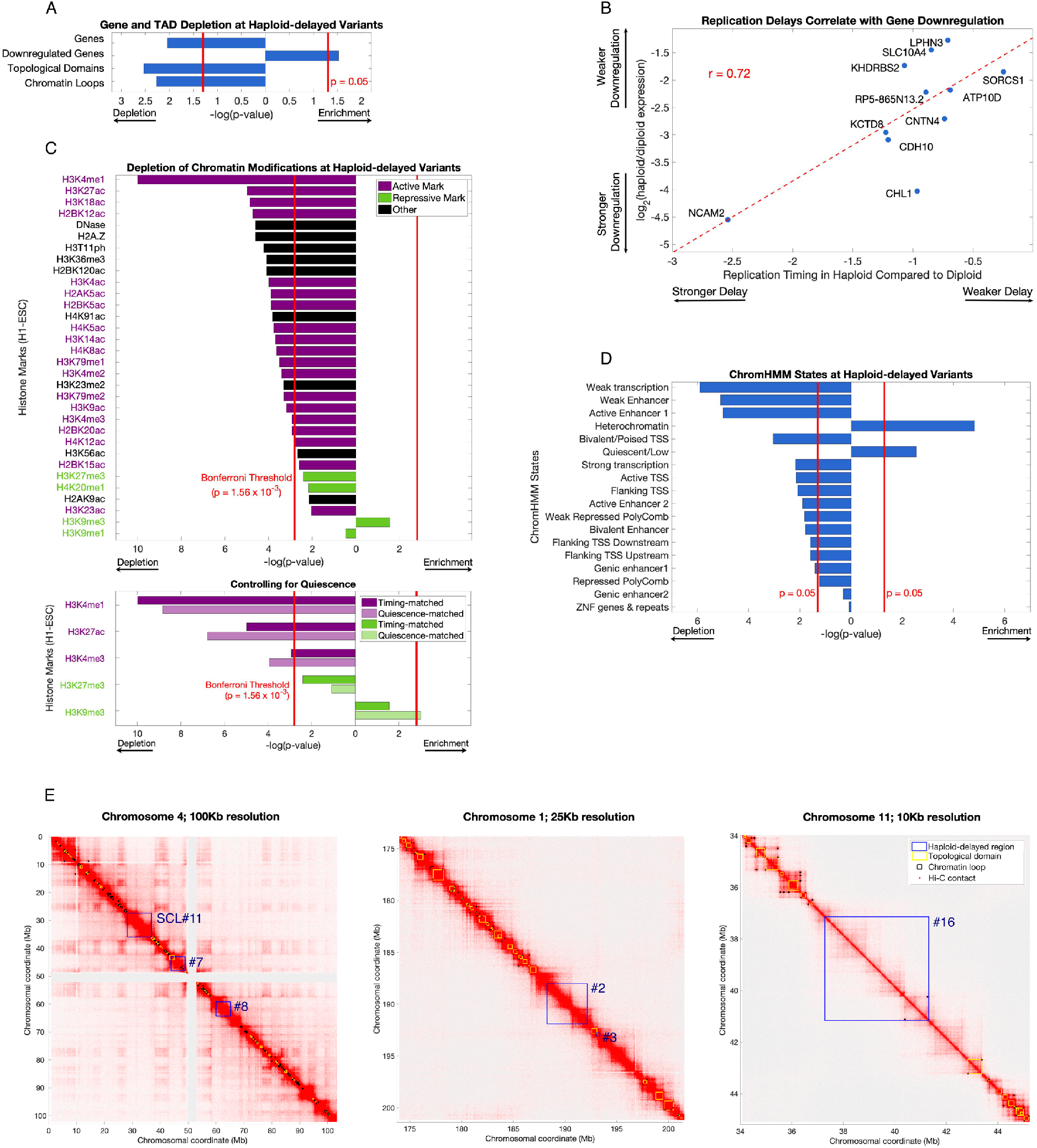
Haploid-delayed variants are located in regions of unorganized, quiescent heterochromatin depleted of genes, histone marks, and 3D contacts. **A.** Haploid-delayed variants show a significant depletion of genes, topologically-associated domains (TADs), and chromatin loops, and an enrichment of genes downregulated in haploid ESCs. Bars indicate enrichment/depletion two-sided t-test p-value of each tested feature in replication timing variants compared to 1000 matched permutations. **B.** The extent of haploid replication delay correlates with the magnitude of gene expression differences between haploid and diploid cells. Spearman rank correlation was similar (rho = 0.65), thus this correlation is not driven by NCAM2. **C.** Haploid-delayed variants are depleted of chromatin marks. Bars shown as in **A**. Each chromatin mark was considered to be independent and the Bonferroni corrected p-value 0.0016 (0.05/32) is shown as a solid red line. Histone mark activation/repression information was obtained from (Schiltz et al., 1999; Zhao and Garcia, 2015). Bottom: three of five tested histone marks (chosen due to their similarity with UR regions) remain depleted when controlling for quiescence, while H3K27me3 rermained nominally depleted, and H3K9me3 became significantly enriched. P-values for matched permutations (dark bars) are identical to the upper panel, with p-values for corresponding quiescence-matched permutations (light bars) shown below them. **D.** As in A and C, for ChromHMM states. Overlap was calculated by the number of basepairs of each state at variant regions. ChromHMM states are mutually exclusive (i.e. not independent), thus a p-value cutoff of 0.05 was used for significance. **E.** Hi-C chromosome interaction maps in H1-ESC (Dekker et al., 2017) at three different genomic locations and resolutions. Haploid-delayed variants (blue) have little overlap with chromosome domains (yellow) or loops (black). This trend is observed across all haploid-delayed variants (panel A).

We next compared the haploid replication timing variant locations to ESC data for 18 ChromHMM states, 31 histone marks, sites of DNase hypersensitivity (Roadmap Epigenomics et al., 2015), and chromosome conformation (topological domains and chromatin loops; (Dekker et al., 2017)). Haploid-delayed variants were significantly enriched for the heterochromatin and quiescent (chromatin lacking histone marks (Hoffman et al., 2013)) states (p = 1.5 × 10^−5^ and 2.8 × 10^−3^ respectivly; **Fig. 3D**) and otherwise significantly depleted for most other chromHMM states. In total, 84.4% of haploid-delayed variant regions were quiescent, while 8.4% were heterochromatic. Of the 32 specific chromatin marks we tested, 31 were nominally depleted in haploid-delayed variants, much more than expected (binomial p = 7.45×10^−9^; **Fig. 3C**). These depletions were statistically significant for 25 of the marks, with a particularly prominent depletion of the active histone mark H3K4me1 (p = 1.04 × 10^−10^). The only nominally enriched mark at haploid-delayed variants was the repressive mark H3K9me3, which is typically associated with constitutive herterochromatin (p = 0.04; not significant after Bonferroni correction).

Haploid-delayed variants were also significantly depleted for both chromosome conformation topological domains (p=0.0037) and chromatin loops (p=0.0067) compared to permutations (**Fig. 3A**). Haploid-delayed variants occupied regions devoid of toploigical domains (**Fig. 3E**). Such regions are stratified based on size into small regions (<50Kb) that constitute topological boundaries and larger regions (>50Kb) that are considered to be “unorganized heterochromatin” (Dixon et al., 2012). The large size (1.62Mb to 14.7Mb) of the regions containing haploid-delayed variants suggests that they all fall into the latter category

Neither replication timing nor the quiescent state explained the observed depletions, as genes (p = 0.033), toplogical domains (p = 0.028), and chromatin loops (p = 0.029) remained significantly depleted from haploid-delayed variants after controlling for quiescence (**Methods**), and downregulated genes remained enriched (p = 0.002). Histone mark enrichment patters were also independent of the quiescent state (**Fig. 3C**). In summary, haploid-delayed variants are late-replicating, depleted of genes, devoid of almost all chromatin marks, and show reduced chromatin contacts and generally unorganized heterochromatin. They thus represent a set of genomic locations with a strong replication phenotype yet are distinct from previously characterized genomic fragile sites.

### Haploid Replication delays correspond to sites of DNA under-replication in mouse placenta polyploid cells

We showed above that a prominent replication aberration in haploid cells is severe delays at 21 regions throughout the autosomes. An attractive possibility is that the severity of these delays, which extends DNA replication well beyond the normal bounds of S-phase, may be related to the frequent diploidization of haploid cells. Furthermore, it is possible that similar replication abberations occur during physiological polyploidization. To begin to test this, we considered analogous instances of replication abnormalities in polyploid tissues. In particular, trophoblast giant cells (TGCs) of the mammalian placenta undergo successive rounds of genome duplication, giving rise to cells with ploidies as high as 1000N (Zybina and Zybina, 1996). These cells do not exhibit uniform DNA copy number, instead a study in mouse TGCs found large genomic regions with reduced DNA copy number in polyploid cells that was thought to be due to DNA under-replication during the multiple replication cycles (to our knowledge, equivalent studies in human placental cells have not been carried out). Under-represented (UR) regions gradually accumulate over time during TGC development, becoming both larger and more numerous in successive cell cycles. Similar to the delayed variants in haploid ESCs, UR regions are late-replicating regions in trophoblast stem cells (Hannibal et al., 2014).

As DNA replication timing is largely conserved between human and mouse (Ryba et al., 2010; Yaffe et al., 2010), we compared 45 UR regions found across mouse TGC development to the 21 regions with replication timing delays in haploid human ESCs (**Methods**). Intriguingly, 11 UR regions each overlapped a separate haploid-delayed variant (**Fig. 1C, Fig. S1C**). When compared to matched random permutations (which maintained the number, size, and replication timing of the haploid-delayed variants; **Methods**), this overlap was highly significant (p = 1.2 × 10^−5^ permuting haploid-delayed variants; p= 4.0 × 10^−3^ controlling for quiescence; p = 1.4 × 10^−9^ permuting UR regions; **Fig. 4A**). The similarity between mouse TGC UR regions and haploid-delayed variants was even more substantial considering that the 11 UR regions corresponded to 13 of the most significant (p < 10^−30^) haploid-delayed variants (**Fig. 1C, Fig. S1C**). In addition, when considering the same UR regions but using the broader genomic coordinates present at late stages of TGC development (Hannibal et al., 2014), we found a substantially increased span of overlap: six of the 11 haploid-delayed variants showed complete overlap with a UR region, while the other five showed substantial (> 48%) overlap along their lengths. This amounted to 64% of the total length of haploid-delayed variants showing co-occurance with UR regions (**Fig. 4B**). The most significant single cell line variant (SCL#11) also encompassed an entire UR region (**Fig. 1C**). Mouse UR regions did not significantly overlap haploid-advanced variants (p = 0.94), however the strongest haploid-advanced variant (#30; **Fig. S1D**) showed near complete correspondence to a UR region (100% overlapped by UR region, and 91% coverage of the UR region). Thus, there is a compelling correspondence of mouse TGC UR regions and haploid ESC-delayed variants, despite the different cell types and species.

**Figure 4.**
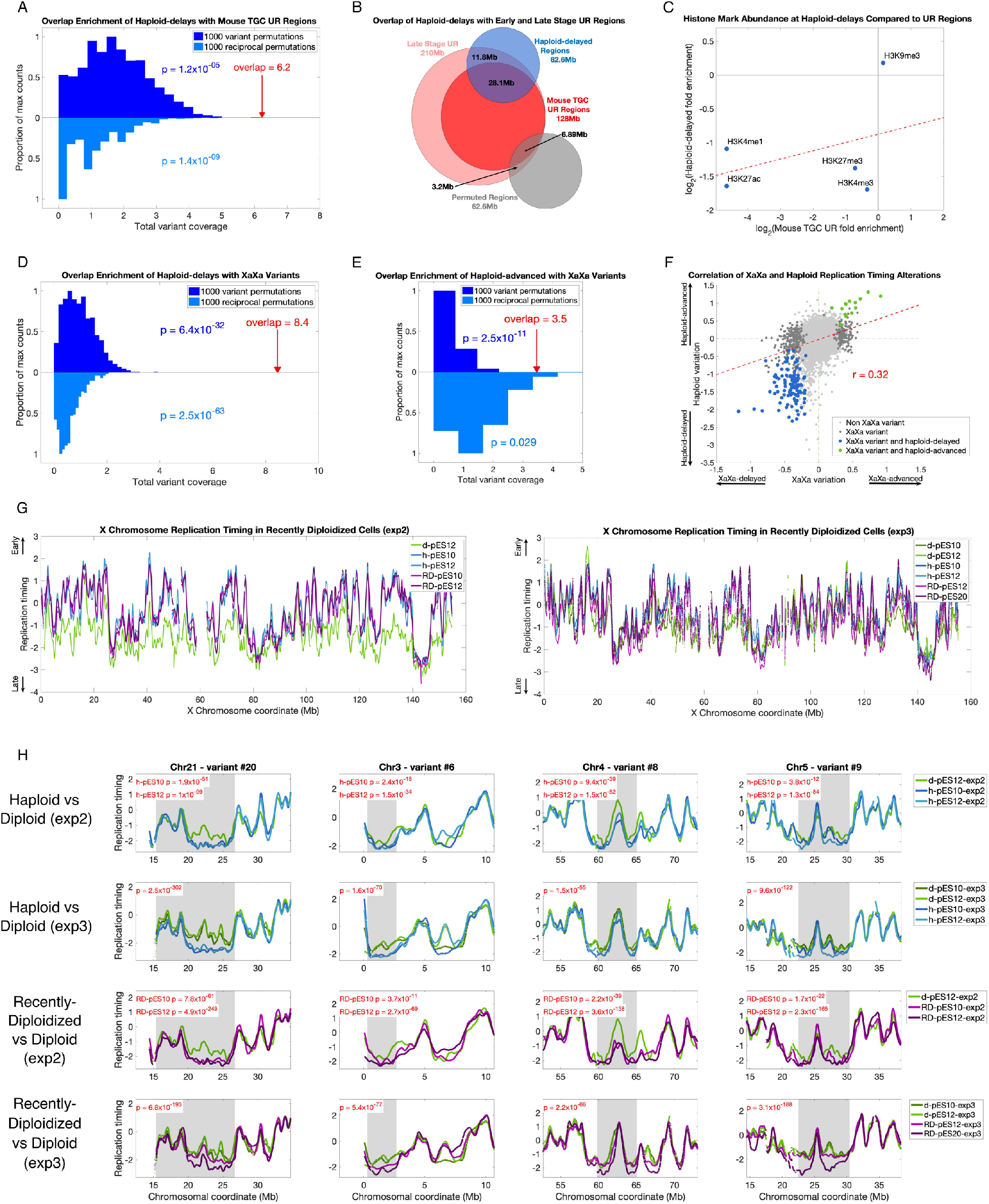
Regions of replication delay in haploid cells correspond to underreplication in polypoid cells and are correlated with the level of X chromosome activity. **A.** Haploid-delayed variants significantly overlap UR regions in mouse TGCs compared to matched permutations. Top histogram (dark blue): permutations of haploid-delayed variants; bottom histogram (light blue): permutations of UR regions. Red arrow: overlaps in actual haploid-delayed variants. Each overlap was counted as the fraction of the haploid-delayed variant that overlapped a UR region (i.e. a value between 0 and 1 for each variant, with a maximum overlap value of 21 corresponding to the 21 haploid-delayed variants). **B.** Venn diagram comparing haploid-delayed variants (blue) to mouse UR regions (red). Dark red: shared regions found in all mouse placenta; light red: regions only identified at later stages of development. Grey: randomly selected permutation. **C.** Histone mark enrichments at haploid-delays resemble those in mouse TGCs at UR regions. **D.** Haploid-delayed variants significantly overlap XaXa variant regions compared to matched permutations. Same as in **A**; light blue: permutations of XaXa variant regions **E.** Haploid-advanced variants significantly overlap XaXa variant regions. Same as **D**, for haploid-advanced variants. **F.** Replication timing differences between XaXa females and other ESCs (x-axis) compared to haploid versus diploid replication timing variants (y-axis). For presentation purposes, data was down-sampled to one window every 200Kb of uniquely alignable sequence. **G.** X chromosome replication timing profiles for haploid, recently-diploidized and diploid ESCs. Haploid and recently diploidized cell lines show comparable replication timing along the X chromosome that is notably earlier than diploid cells. This trend is more pronounced in experiment 2 (left) than in experiment 3 (right). **H.** Replication timing in recently diploidized ESCs resembles haploids more than it resemble diploids, thus tracking X chromosome status rather than ploidy. For experiment 2, p values (t-test) are shown for each cell line compared to the single diploid cell line in that experiment; for experiment 3, a single p value (ANOVA) is shown for the joint comparison of the two haploid cell lines with the two diploid cell lines. See **Table S2** for statistics at other haploid-delayed variants.

Strikingly, UR regions in mouse TGCs are depleted for H3K4me1, H3K27ac, H3K4me3 and H3K27me3 and enriched for H3K9me3 (Hannibal et al., 2014) – the exact same trends observed for haploid-delayed variants (**Fig. 3C**). Furthermore, there was a correlation in the level of enrichment of these five histone marks between haploid-delayed and mouse UR regions (**Fig. 4C**). In further similarity to haploid-delayed variants (**Fig 3A**), UR regions were depleted of genes compared to the rest of the genome (Hannibal et al., 2014). The genes that were found in mouse UR regions were enriched for cell adhesion and neurogenesis annotations, similar to haploid-delayed variants, which contained both cell-adhesion genes and several categories of genes related to the nervous system (GABA-A receptor activity, neuron part, and postsynaptic membrane annotations) (**Table S1**).

Taken together, there is a strong correspondence of mouse polyploid TGC underreplicated regions and haploid-delayed variants in terms of genomic location, replication timing, histone composition, and gene content. These similarities could represent common biological mechanisms linked to whole genome duplication. A key difference is that delayed replication in haploid cells cannot possibly be a consequence of diploidization, since it precedes it.

### Replication delay in haploid cells is linked to X chromosome dosage

We next considered possible mechanisms leading to the replication defects in haploid cells. In particular, we considered whether the relative increase in X-linked gene expression in haploid cells (Sagi et al., 2016) could influence autosomal replication timing and account for the replication delays we observed. To test this, we first sought to find other instances of increased X chromosome expression dosage in ESCs. Female ESCs occasionally undergo partial or complete reactivation of the inactive X chromosome (Patel et al., 2017). We therefore searched for cell lines with evidence of a reactivated X-chromosome by utilizing the replication profiles of 116 human ESCs (Ding et al., 2020). While replication timing was highly correlated among cell lines along the autosomes (mean r = 0.9) as well as the X chromosome in male (karyotypically XY) samples (r = 0.91), it was significantly less correlated on the X chromosome of female samples (mean r = 0.83; **Fig. S3A**), as expected due to the random replication of the inactive X chromosome (Koren and McCarroll, 2014) (similar patterns were observed when comparing haploid and diploid cells, see **Fig. S1B**). However, we identified nine female samples with X chromosome replication timing patterns that resembled male more than they resembled other female cell lines. These cell lines were identified by their high correlations to male samples on the X chromosome (**Fig. S3B;** those with r > 0.89), yet also showed higher correlations among them (0.86-0.92; similar to autosomes or the X chromosome in male samples) and lower correlations to other female samples (0.78-0.84). Furthermore, the X chromosomes of these cell lines replicated earlier than other female samples (−0.30 compared to −0.87; **Fig. S3C**). We suspect that these female cell lines may have undergone partial or complete reactivation of the inactive X chromosome.

When comparing the nine cell lines with suspected X chromosome reactivation (designated “XaXa”) to the other 107 ESCs, we identified 214 autosomal regions with subtle, yet significant replication timing variation between the two (**Methods**). This suggests that the dosage of active X chromosomes may be linked to replication timing alterations genome-wide. Strikingly, 36 of the 214 reactivated-X variants overlapped 16 of the 21 haploid-delayed variants (**Fig. 1C**). This represents a highly significant enrichment compared to expectation (p = 6.4 × 10^− 32^ when permuting haploid-delayed variants; p = 6.4 × 10^−33^ when controlling for quiescence; p = 2.5 × 10^−63^ when permuting reactivated-X variants; **Fig. 4D**). Furthermore, six reactivated-X variants overlapped five different haploid-advanced variants, again much more than expected by chance (p = 2.5 × 10^−11^ and p = 0.029 when permuting haploid-advanced variants or reactivated-X variants, respectively; **Fig. 4E**). All of the 42 overlapping regions showed a consistent direction of replication timing change in XaXa ESCs and in haploid ESCs, such that when haploids were delayed (or advanced) relative to diploids, XaXa ESCs were also delayed (or advanced) relative to XaXi ESCs. Across all these genomic regions, XaXa and haploid replication timing differences were well correlated (r = 0.32; **Fig. 4F**). These results are consistent with our premise by which replication timing alterations in haploid cells could be related to the elevated dosage of X chromosome gene expression.

To more directly test whether X chromosome activity could be causing the replication timing alterations in haploid cells, we profiled replication timing in recently diploidized (RD; haploid cell cultures shortly after becoming diploid) alongside haploid and diploid (XaXi) cells. X chromosome inactivation occurs only several cell divisions after cells become diploid, thus shortly after diploidization, the two X chromosomes are both active (Sagi et al., 2016). This was supported by our data, as X-chromosome replication timing in recently diploidized cells was similar to haploid cells and much earlier than in later-stage diploid cells, consistent with the presence of two active X-chromosomes (**Fig. 4G**).

Across two separate experiments (experiments “2” and “3”), we reproduced between 15 and 18 (71-86%, three delays only validated in h-pES12) of the 21 variants in experiment 2, and 18 (86%) in experiment 3 when comparing haploid and stably diploid cells (**Fig. 4H; Table S2**). Notably, the replication delays in these repeat experiments were dampened compared to the original experiment; we suspect that this is related to “culture adaptation”, in which late-passage, serially-sorted haploid cell lines become more stable in the haploid state and undergo reduced rates of diploidization.

Across the majority of variant regions, RD cell lines showed the same replication delays as haploid cells and were distinct from stably diploid cells. Of the 21 haploid-delayed variant regions, between 14 and 19 (67-90%; experiment 2, five variants found only in RD-pES12) and 19 (90%; experiment 3) were also significantly delayed in RD cell lines compared to later-stage diploid cell lines, despite both being diploid (**Fig. 4H; Table S2**). In particular, of the 15 regions delayed in all haploid cells across all three experiments, all were also delayed in all RD cell lines. Interestingly, one of the RD cell lines (RD-pES12-exp3) showed comparatively weaker delays than other RD cells (**Fig. 4H**), and also showed X-chromosome replication timing more consistent with later-stage diploids; this may indicate that this cell line partially inactivated one X chromosome, which is consistent with it having been cultured for many (34) passages. Taken together, haploid-delays correlate more strongly with the dosage of active X chromosomes than with ploidy *per se*. This provides compelling support to the notion that X chromosome dosage could underlie the DNA replication delays in haploid ESCs.

Finally, the repeat experiments and inclusion of recently-diploidized cell lines further clarified the haploid-diploid differences on the X chromosome. Specifically, while the X chromosome generally replicated earlier in haploid compared to diploid cells, we noticed two regions that seemed not to exert this difference (**Fig. 1A**). While this observation in the initial experiment was suspected to result from technical noise, these regions reproducibly showed this same effect in both repeat experiments as well as in the RD cell lines (**Fig. S3D**). We thus propose that these are likely regions of haploid replication delays on the X chromosome. This possibility is also consistent with these two regions being long and late-replicating, similar to autosomal haploid-delayed regions. Thus, we suggest that there are a total of 23 regions across the genome identified here as having delayed replication in haploid ESCs.

## Discussion

Haploid human stem cells provide a powerful system for human genetic studies as well as a unique opportunity for investigating the biology of cell ploidy. However, their spontaneous diploidization is a major limitation to their use and remains a poorly understood phenomenon. Here we show that DNA replication is delayed well beyond the bounds of S-phase at multiple regions in human haploid embryonic stem cells. The replication delays we describe are among the most profound alterations to the otherwise highly stable eukaryotic DNA replication timing program. The ability to identify replication delays that transcend S phase was enabled by our approach of directly sequencing DNA from proliferating cell cultures; previous replication profiling approaches that utilize FACS sorting of S phase cells (Hulke et al., 2020) would likely have missed these extreme delays as G2 cells are not included, and thereby the valleys of very late replicating regions are not fully captured.

Haploid-delayed regions have a distinctive genetic and epigenetic signature, characterized by late replication, a paucity of genes, limited histone modifications, and reduced chromatin contacts. They differ from previously described chromosomal fragile sites. In contrast, they show profound similarity to UR regions in mouse placenta trophoblast giant cells. Thus, these regions may represent a novel class of co-regulated genomic sites that are susceptible to abnormal replication and linked to polyploidization in the placenta, and potentially other cell types. Finally, we provide compelling evidence implicating the dosage of X chromosome activity, rather than the haploid state *per se*, in replication delays. Similar replication delays were observed in ESCs with evidence of a reactivated X-chromosome and, more importantly, in diploidized ESCs with two active X chromosomes.

### What causes replication delays in haploid cells?

We can envision two mechanisms by which X chromosomes could cause replication delays. First, the mere presence of an inactive X chromosome may be the critical factor. An inactive X chromosome may recruit heterochromatin factors, sequestering them from other chromosomes. An absence of an inactive X, on the other hand, may release such factors to bind to specific autosomal regions and delay their replication. This is similar to the “chromatin sink” model suggested for the highly heterochromatic *Drosophila* Y chromosome (Francisco and Lemos, 2014). This hypothesis can be tested with human cell lines lacking an inactive X chromosome, such as Turner syndrome (X0) cells. We profiled replication timing in an XX H9 ESC line and an X0 progeny that was cloned after spontaneous loss of one of the X chromosomes. We only identified four of the 21 haploid-delayed regions (as well SCL#11) as delayed in X0 compared to XX ESCs(**Fig. S3E**). While notable, the lack of more supstantial corrrepsondence between X0 delays and haploid-delayed regions suggests that the absence of an Xi is insufficient to explain the full complement of haploid replication delays.

An alternative hypothesis is that overexpression of an X-linked gene(s) contributes to the observed replication delays. Thus, a single active X chromosome in haploid cells, or two active X chromosomes in diploid cells, could produce an elevated level of a transcript(s) compared to normal X_a_X_i_ diploid cells. This overexpression could then potentially cause replication delays at specific loci across the genome. Several X-linked genes are possible candidates for mediating autosomal replication timing delays. For instance, ELK1 is an X-linked transcription factor primarily expressed in placenta and ovarian tissue (Uhlen et al., 2015) that was shown to cause transcriptional changes of autosomal genes in reactivated-X ESCs (Bruck et al., 2013); PPP2R3B encodes a protein phosphatase 2A subunit that delays the firing of replication origins throughout the genome by stabilizing the Cdc6-Cdt1 interaction (van Kempen et al., 2016); BCOR, a Polycomb-group repressive complex gene, is required for normal placental development (Hamline et al., 2020); and NAP1L2, which encodes a nucleosome assembly protein, is induced during differentiation of mouse trophoblast stem cells to TGCs (Attia et al., 2007) and is upregulated in human haploid ESCs (Sagi et al., 2016). The role of these and other genes in haploid replication delays could be tested by knockout, knockdown or overexpression in haploid and diploid ESCs.

### Can delayed replication cause diploidization?

The mechanisms causing haploid cells to diploidize, or diploid cells to become polyploid, are not fully understood. It is intriguing to consider whether the severe replication delays we described in haploid ESCs could induce diploidization, and whether similar mechanisms could be operative in other instances of polyploidization. It is known that incomplete DNA replication leads to an ATR-dependent activation of the S-M checkpoint that prevents cells from prematurely entering mitosis (Enoch et al., 1992; Eykelenboom et al., 2013). Thus, it is possible that replication delays ultimately lead haploid cells to avert mitosis and re-enter the cell cycle, making them diploid. In support of this possibility, it was previously shown that in human cancer cell lines, chromosome-scale replication delays and associated delays in mitotic chromosome condensation activate the S-M checkpoint and result in endoreplication in a subset of cells (Chang et al., 2007). Disruption of the replication initiation factors ORC2 or GINS2 has also been shown to induce polyploidization in human cells (Huang et al., 2016; Rantala et al., 2010). Similarly, it was recently shown that ~2% of the maize genome is delayed in its replication in endocycles compared to normal mitotic cycles (Wear et al., 2020). Polyploidization is also very common in cancer (Bielski et al., 2018), and so is replication stress; it is intriguing to consider whether these two phenomena are related to each other. To further test these links, we induced replication delays in haploid ESCs using Aphidicolin, a DNA polymerase inhibitor. Over three independent experments, we observed a significant increase in diploidization rates 48hrs after Aphidicolin treatment, consistent with incomplete replication promoting diploidization of these cells (**Fig. S3F**). Another observation supporting a general restructuring of the cell cycle is that not every chromosome contained regions of replication delay in haploid cells. This suggests that diploidization is not due to replication delays directly causing mitotic chromosome segregation defects on the chromosomes on which they occur, but rather that one or more delayed regions can activate a global cellular response (e.g. a checkpoint) that could lead to diploidization.

The fate of un-replicated DNA during mitotic entry has been studied before, for instance in the context of common fragile sites following replication stress. A mechanism, termed mitotic DNA synthesis (MiDAS), has been described in which DNA repair synthesis is initiated at chromosomal gaps or breaks during mitosis. MiDAS utilizes a pathway resembling RAD52-dependent break induced replication (BIR; (Bhowmick et al., 2016; Minocherhomji et al., 2015)). This raises the possibility that lack of homologous chromosomes renders haploid cells particularly vulnerable to incomplete replication since they are unable to perform MiDAS. Delayed replication in haploid cells would thus induce a robust checkpoint response. In contrast, diploid cells don’t suffer the replication delays that require replication completion in mitosis, while recently diploid cells have replication delays similar to haploids but are competent at MiDAS given the presence of homologous chromosomes. Indeed, we do not observe frequent tetraploidization of recently diploid cells despite them having replication delays. This model would also imply that haploid cells are able to eventually complete genome replication and/or repair any associated DNA damage following diploidization. Consistently, we do not observe any gross or recurrent genome rearrangements in diploidized cells. Such repair synthesis could potentially occur in 53BP1 nuclear bodies during G1 phase following diploidization (Bhowmick et al., 2016; Harrigan et al., 2011; Lukas et al., 2011; Minocherhomji et al., 2015).

In support of the idea that haploid-delays are linked to whole genome duplication, we observed a remarkable correspondence between these delays and UR regions in mouse TGCs. Haploid delays and URs are found in corresponding genomic locations, have similar replication timing, contain nearly identical histone patterns, and are enriched for some of the same gene categories. The replication factor Rif1, which is required for underreplication in *Drosophila* (Munden et al., 2018), was also suggested to be important for under-replication in mouse TGCs (Hannibal and Baker, 2016). Furthermore, the inactive X chromosome in mouse TGCs is unusual in harboring both the heterochromatic mark H3K27me3 and euchromatic marks such as H3K4me2 (Corbel et al., 2013) and a high fraction of genes that escape X chromosome inactivation (Schoenfelder and Fox, 2015). However, in male embryo pregnancies placental cells only carry a single X chromosome, thus the mechanism of polyploidization in TGCs is likely independent of X chromosome dosage, although may still be related to a similar gene circuitry as the one putatively disrupted in human haploid ESCs. Mouse UR regions were originally identified using microarray genomic DNA hybridization (Hannibal et al., 2014). However, TGC profiling using next-generation sequencing suggest that DNA copy number decreases gradually rather than sharply at UR regions (see (Hannibal et al., 2014) Figure 2; (Hannibal and Baker, 2016) Figure 1). This, and the absence of evidence for chromosomal deletions at UR regions using paired-end sequencing (Hannibal et al., 2014) may be more consistent with the interpretation that UR regions represent severely delayed replication, similar to haploid delays, rather than DNA loss.

Further work is required in order to understand the interplay between X chromosome dosage, transcriptome remodelling, replication dynamics, and whole genome duplication. Key remaining questions are why haploid cells become diploid at high rates whereas the diploid state is much more stable; and whether similar replication-related mechanisms, whether dependent on X chromosome activity or not, contribute to polyploidization in TGCs and other tissues. Extending our analysis to mouse haploid cells, androgenetic embryonic stem cells, differentiated cells, and genetically-manipulated cells will shed light on the fundamental links between genome regulation and ploidy control, and could ultimately enable the stabilization of the haploid state in human ESCs.

Our results suggest that replication dynamics in S phase have the potential to influence the entire cell cycle. It is thus critical for cells to maintain their temporal order of DNA replication. As a corollary, certain DNA sequences can have a physiological role by virtue of their replication properties rather than their actual coding potential. This could ascribe a development function to late DNA replication, in particular by various mammalian cell types (Sagi and Benvenisty, 2017) exploiting site-specific DNA replication delays in order to become polyploid.

---

**Supplementary Figure 1.**
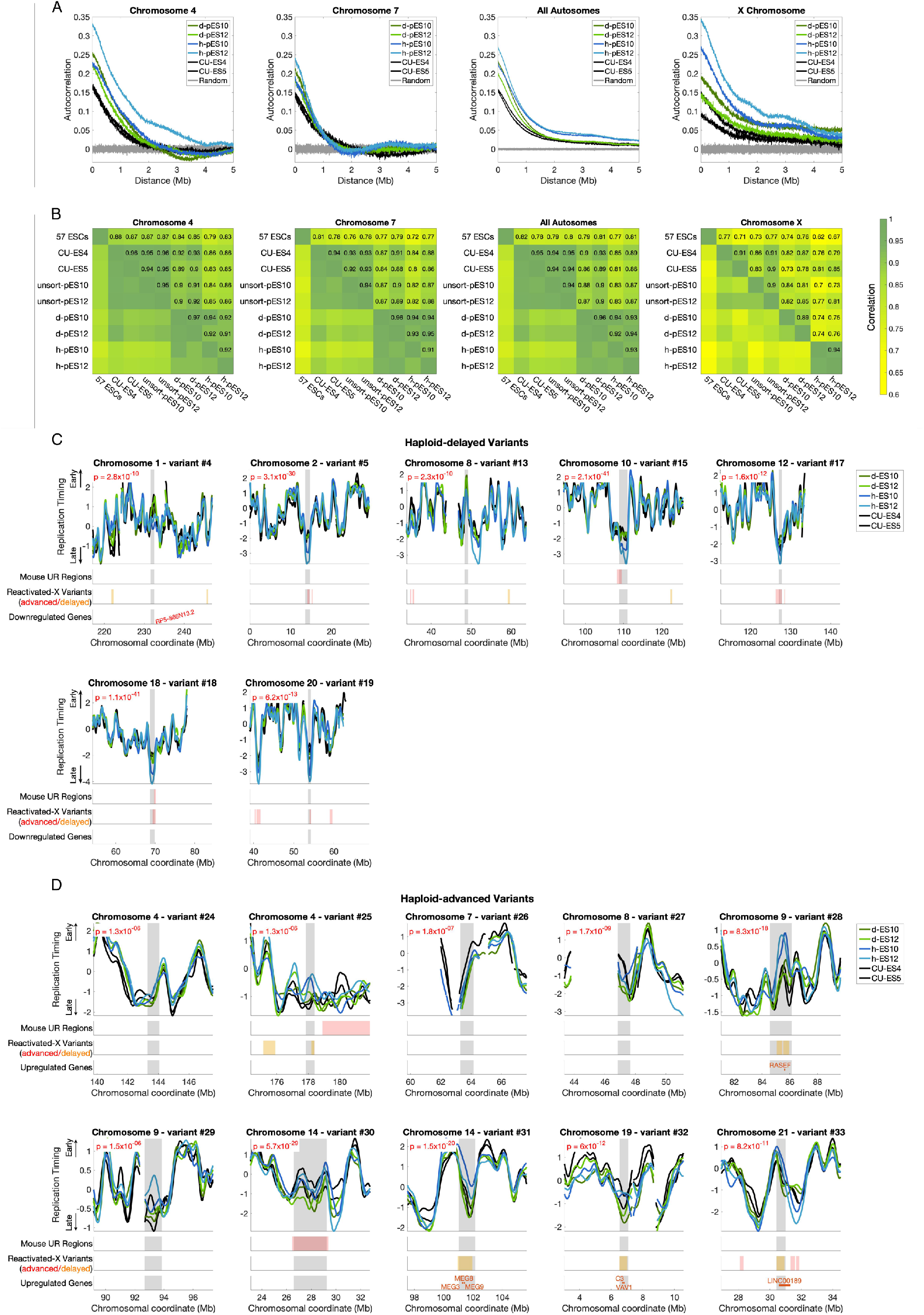
Data QC and additional replication timing variants (not shown in Fig. 1). **A.** Autocorrelation of raw (unsmoothed) DNA replication timing profiles across two sample chromosomes (4 and 7), all autosomes combined, and the X chromosome, for haploid, diploid, and control cell lines. **B.** Pearson correlations of smoothed replication timing profiles across the same chromosomes as in A. “57 ESCs”: mean replication timing of 57 previously measured female ESCs with evidence of normal X chromosome inactivation. unsort: unsorted for ploidy. **C.** All additonal haploid-delayed variants ordered by genomic position and plotted as in **Fig. 1C**. **D.** Haploid-advanced vaiant regions, ordered by genomic position. Note that variant #30 appears to be a diploid-delay rather than a haploid-advance, as diploidized ESCs show later replication than both haploid and the two control ESCs. Upregulated rather than downregulated genes are shown within the variant region.

**Supplementary Figure 2.**
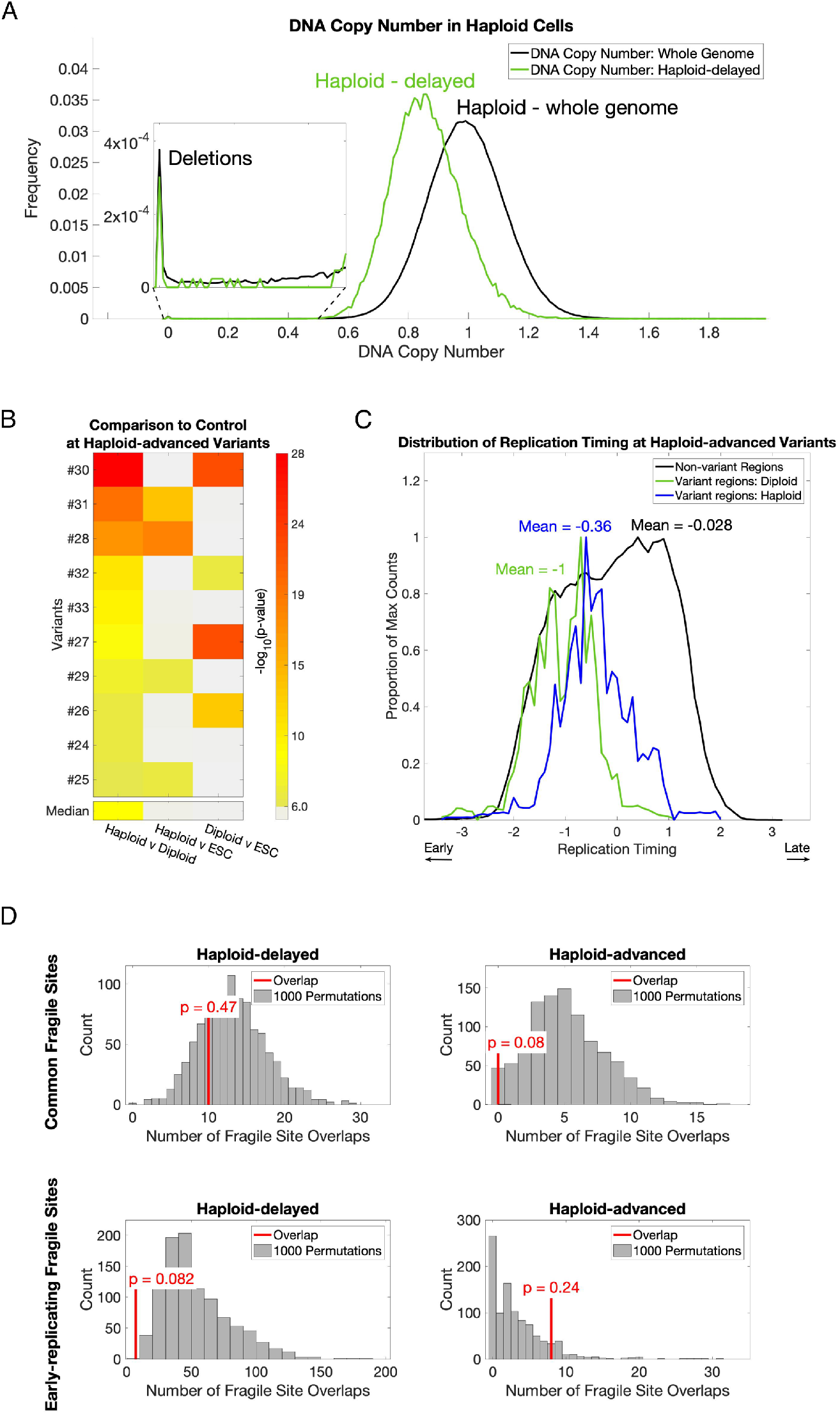
Further characterization of haploid-delayed and haploid-advanced regions. **A.** Haploid cell line DNA copy number at all genomic loci (black) and at haploid-delayed variants only (green). Loci with a copy number near zero (inset) are considered to represent deletions. They do not correspond to unmappable loci as loci with no reads in neither haploid, diploid, nor control (CU-ES4 / CU-ES5) cell lines were removed from further analysis. Thus, haploid-delayed variants are not attributed to chromosomal deletions (see further in **Methods**). **B.** Same as Figure 2D, for haploid-advanced variants. Four variants (#31, #28, #29 and #25) show significant variation between haploid cells and control ESCs, while four variants (#30, #32, #27, and #26) show significant variation between diploid cells and control ESCs (two were ambiguous). **C.** Same as Figure 2E, for haploid-advanced regions. Haploid-advanced variants on average replicate relatively late in diploids and closer to mid-S phase in haploids. **D.** Overlap of haploid-delayed variants (left) and haploid-advanced variants (right) with common fragile sites (CFS) (Bignell et al., 2010; Fungtammasan et al., 2012; Savelyeva and Brueckner, 2014) (top) and early-replicating fragile sites (ERFS) (Barlow et al., 2013) (bottom) compared to 1000 matched permutations (**Methods**). The number of overlaps are shown as red lines with associated p-values. Rare fragile sites (RFS) (Bignell et al., 2010; Zlotorynski et al., 2003) also did not have overlaps with neither class of variants (not shown).

**Supplemental Figure 3.**
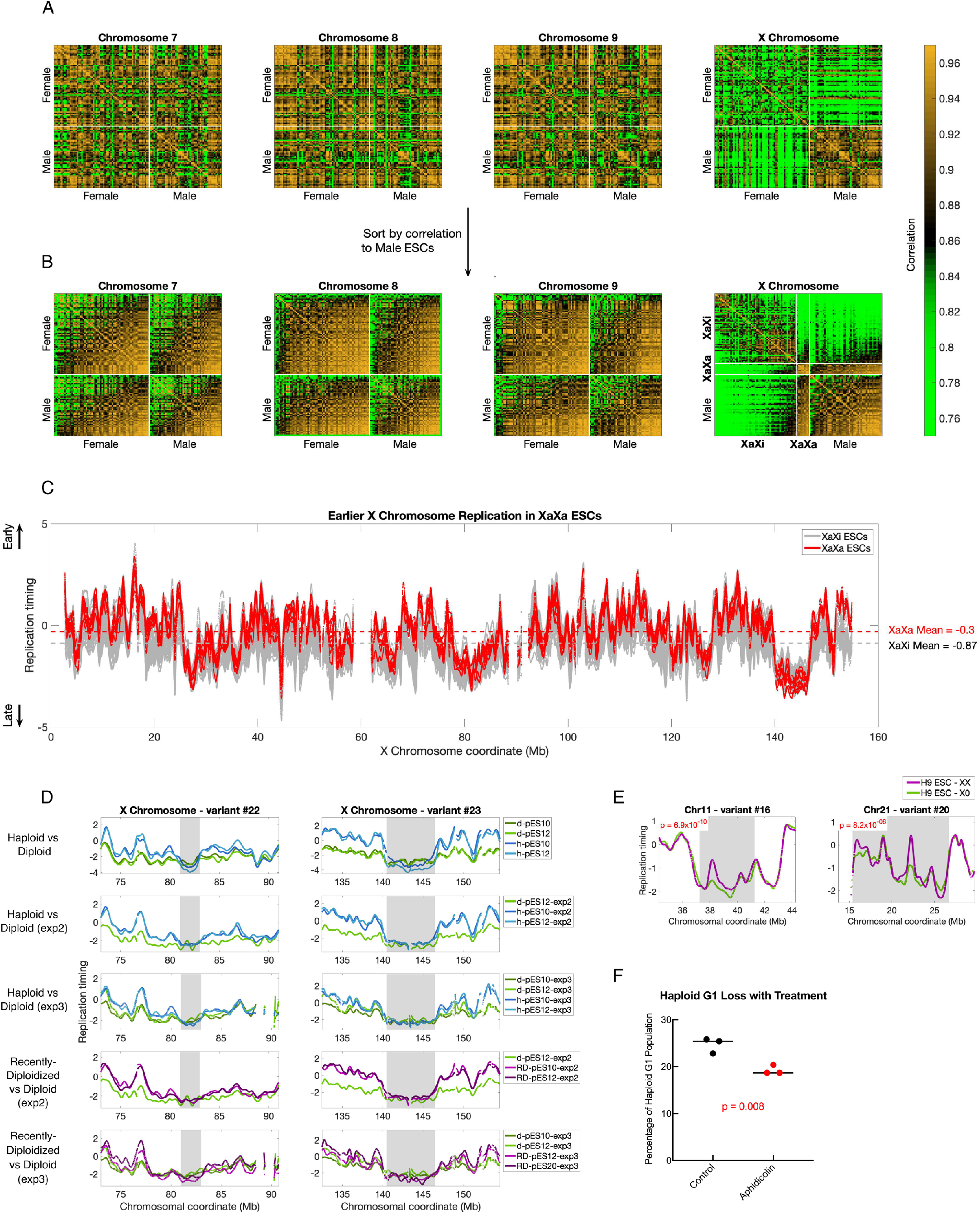
Identification of female ESCs with evidence of X-chromosome reactivation. **A.** Correlation matrices of replication timing across 116 ESCs on chromosomes 7, 8, 9, and X, sorted by sex. **B.** Same correlation matrices as in A, sorted first by sex, and then by mean correlation to male ESCs on each chromosome. Nine female ESCs (marked by white bars) have higher correlations to each other and to male ESCs than to other female ESCs. **C.** X chromosome replication timing profiles for nine XaXa lines (red) and 57 XaXi lines (grey), showing the earlier mean replication timing of XaXa cell lines compared to XaXi cell lines. **D.** The two X chromosome regions are consistently similar between haploid, diploid and recently diploid cell lines, which contrasts them with the remainder of the chromosome replicating earlier in haploid compared to diploid cells. This consistent deviation from the chromosome trend nominates these two regions as additional putative haploid-delayed variants. **E.** The two strongest regions of X0 delay at haploid delayed variants. P-values (t-test) between H9 X0 and H9 XX were calculated across haploid-delayed regions. Seventeen of 21 haploid-delayed regions showed nominally later replication timing in X0 compared to XX, of which four were significantly delayed (Bonferroni p-value = 2.4 × 10^−3^). Variant SCL#11 was also delayed in H9 X0 (not shown). **F.** Diploidization following replication delays induced by Aphidicolin. Shown are the percentage of haploid G1 cells, evaluated using flow cytometry, after 48 hrs of treatment with DMSO (control) or 0.3mM aphidicolin, followed by an additional seven days of culturing without drug. P-value was determined using a t-test.

## Acknowledgments

N.B. is the Herbert Cohn Chair in Cancer Research. This work was partially supported by the Israel Science Foundation (grant no. 494/17), by the Rosetrees Trust, and by the Azrieli Foundation (to N.B.), by NYSTEM IDEA award #C32564GG (to D.E.), and startup funds from Cornell University (to A.K.).

## Data Availability Statement

Data of hESC lines sequenced in this study were desposited in dbGaP (accession number: phs001957).

## Methods

### Cell culture

Haploid pESCs, diploid pESCs, and control ES cell lines were cultured and maintained as previously described (Sagi et al., 2016). Briefly, we used StemFlex media on Geltrex matrix at 37°C with 5% CO_2_ and atmospheric oxygen concentration. Passaging was performed with trypsination via incubation with TryplE and culturing newly passaged cells with 10uM ROCK inhibitor (Y-27632) for 24 hours. Freezing was performed with media consisting of 10% DMSO and 40% FBS. Haploid and diploid cell lines were FACS-sorted in G1 phase, and plated and cultured for 3-7 days in order to ensure the desired ploidy as well as the cell cycle asynchrony of the cultures (which is a pre-requisite for the replication timing assay).

### Whole genome sequencing

DNA was extracted with the MasterPure DNA purification kit (Lucigen). Libraries were prepared using the Illumina TruSeq PCR-free library preparation kit and sequencing was performed on the Illumina NextSeq500 with 75bp single-end reads (first experiment), or the Illumina HiSeq X Ten with 150bp paired-end reads (second and third experiments). Reads were then aligned to the hg19 human reference genome using BWA-MEM (Li and Durbin, 2010).

In the first experiment, haploid and diploid pES10 and pES12 were sequenced alongside three control ESCs. One control ESC (B123) showed poorer correlations and autocorrelations than other samples and was removed from further analysis.

In the second experiment, h-pES10, h-pES12, RD-pES10, RD-pES12, and d-pES12 were sequenced. In the third experiment, we again sequenced h-pES10, h-pES12, d-pES10, d-pES12 (two separate samples), as well as RD-pES12 and RD-pES20. Importantly, RD-pES12 was sequenced at passage 9, while RD-pES20 was sequenced at 34 passages. One of the two d-pES12 samples was removed from further analysis due to poor data quality

### Generation of DNA replication timing profiles

In order to infer replication timing, we first used GenomeSTRiP to infer DNA copy number across the genome (Handsaker et al., 2015; Koren et al., 2014). Sequence read depth was calculated in 2.5Kb windows along the genome, corrected for alignability and GC content. Copy number values for both haploid and diploid cells were normalized to an average DNA copy number of two. These copy number values were then filtered as follows:

1. Windows spanning gaps in the reference genome were removed.
2. Windows with copy number greater than one above or below the median copy number were removed.
3. In order to remove extreme data points, the data was segmented using the MATLAB function *segment*, which groups consecutive data points into segments based on a tolerance threshold. This analysis was done twice using two different segmentations parameters of 0.5 (less strict) and 0.1 (more strict). By using two different parameters, both shorter and larger genomic regions that deviate from the median can be captured. Segments falling above or below the median by a threshold of 0.45 copies were removed.
4. Genomic regions that were further than 30Kb from other data points, and that were less than 300Kb long were removed.
5. Regions shorter than 100Kb between removed data points were removed.
6. Regions shorter than 500Kb between three or more removed data points were removed.

Data was then smoothed using the MATLAB function *csaps* with smoothing parameter of 10^−17^, and then normalized to a median of 0 and a standard deviation of 1, such that positive values represent early replication and negative values represent late replication.

### Identifying replication timing variations

In order to identify replication timing differences between groups of samples (e.g. haploid and diploid cell lines), we used ANOVA across “regions” tiling each chromosome. ANOVA tests the null hypothesis that all samples come from a population with the same mean versus the alternative that each group (haploid or diploid in this case) is drawn from populations with differing means. ANOVA was applied on filtered, raw (unsmoothed) replication timing values. Both the region size and overlap between adjacent regions were optimized by finding the false discovery rate (FDR) for a given set of parameters. To determine the FDR, ANOVA was repeated comparing permuted samples to disrupt the haploid-diploid comparison, i.e. h-pES10 and d-pES12 were compared to h-pES12 and d-pES10. Because this permuted scan compares neither cell ploidy nor genetic background (pES10 vs pES12), we considered significant regions arising from this test to be false. By dividing the total length of these false regions (after several filtering steps- see further below) by the total size of autosomal variants (after the same filtering steps) found between the haploid and diploid cell lines, we determined an FDR for regions of the chosen length. We chose a region size of 76 replication timing data windows (covering 190Kb of uniquely alignable sequence), with a slide of a quarter region (such that each genomic locus was tested four times) in order to optimize the specificity and sensitivity of variant detection; this resulted in an FDR of 0.07. Using this FDR, we identified 14.2-fold more autosomal variation in the haploid-diploid comparison compared to the permuted sample comparison.

Regions with a Bonferroni-corrected ANOVA p < 9.58 × 10^−7^ (0.05/52,184 regions tested; note that we stringently regard each region as independent) were merged into continuous replication timing variants, of which an initial 57 were identified. Then, we removed variants in which pairwise comparisons of haploid and diploid profiles showed differences in the direction of effect (i.e. one haploid-diploid pair showed earlier haploid replication while another showed earlier diploid replication), trimmed variants in which any pair of a haploid and a diploid cell line had overlapping replication timing profiles (i.e. one pair was not variant in a given region), and removed any variants that, after being trimmed, were shorter than the original tested region size (190Kb) or that fell below the significance p-value threshold. This resulted in 53 autosomal variants. These regions were then extended in both directions as long as all haploid and diploid profiles remained separated (i.e. were still variable in the same direction). Any variants that were overlapping, or were nearby (<750Kb) or separated by small gaps and appeared to result from the same region of variation, were merged. Additonally, one variant region on chromosome 4 that occupied a portion (chr4:30,583,544-31,202,423) of variant SCL#11 was removed, as this variant region was both much smaller and weaker (in terms of both p-value and extent of delay) than the encompassing SCL#11 variation in pES12 alone. This resulted in the final set of 21 haploid-delayed variants and 10 haploid-advanced variants.

In order to identify regions in which only one haploid cell line had different replication timing compared to diploid cells, we performed a genome-wide scan similar to the one above but utilizing a t-test, rather than ANOVA, and comparing one pair of samples at a time (this approach was also employed for experiment 2, as we were only comparing each haploid cell line to a single diploid cell line). In each 190Kb region, replication timing in a given haploid cell line was compared to the mean diploid replication timing in that region. Candidate replication timing variants were filtered as above, with the exception that the filter for consistent direction of effect was no longer applicable with only one haploid sample. In addition, we removed any part of these variants that overlapped the shared variants found using ANOVA (with the exception of SCL#11, in which the smaller nested shared variant was removed instead of the single cell variant (see previous paragraph)). This t-test approach was also used to identify replication timing variants in recently diploidized cells, where we compared the mean of recently diploidized cells to the single d-pES12 sample.

For identification of replication timing variants between the nine reactivated-X individuals and the other 107 individuals among the 116 ESCs, we used 20 instead of 76 windows for the ANOVA tests, since these data were in 10Kb instead of 2.5Kb windows.

In order to test whether haploid-delays, identified in the initial experiment, were also present in the second and third experiments, and to test for the presence of these variations in the recently-diploidized ESCs, we performed ANOVA specifically at the haploid-delayed variant regions. Filtered, unsmoothed replication timing data from haploid and recently-diploidized pESCs were each compared to the concurrently sequenced diploid pESCs in each experiment, and p-values less than 2.4 × 10^−3^ (Bonferonni correction of .05/21 for each variant tested) were considered significant (**Table S2**).

### Testing whether haploid-delayed variants are genomic deletions

Since we detect regions with strongly delayed replication, we considered whether these could be due to deletions of the underlying sequence. Even though our analysis filters for regions with DNA copy number significantly different than a sample’s ploidy, it is still possible that the replication variants are influenced by deletions that passed this filtering. However, several aspects of the data argue against the replication variants being chromosomal deletions. First, we expect deletions in haploid cells to have a copy number near zero. In support of this, we found that 46.2% of loci with copy number below 0.1 overlap deletions identified by the 1000 Genomes Project (Sudmant et al., 2015). In contrast, at haploid-delayed variants, while the absolute copy number was lower than the rest of the genome, it was still much greater than zero (mean of 0.86, compared to a genome average copy number to 1, **Fig. S2C**). Within haploid-delayed variants the frequency of suspected deletions (copy number < 0.1) was extremely small (0.0037%), and the only deletions within haploid-delayed variants (variants #9 and #20) were found in both haploid and the corresponding isogenic diploid cell lines, indicating that they are not ploidy-dependent copy number variations (**Fig. S2C**). These analyses, however, do not rule out the possibility of subclonal deletions, in which only a subset of the cells have a deleted region thus giving rise to a copy number value intermediate between zero and one when analyzing a population of cells. If this was the case, we would expect different sub-clonal deletions to have a range of copy number values between zero and one. In contrast, all haploid-delayed variants were much closer to a copy number of one than they were to zero (**Fig. S2C**). Our finding that the copy number at variant regions is highly consistent between the two haploid cell lines (**Fig. 1D)** also argues against the replication variants being sub-clonal deletions, as it would be extremely unlikely for these to occur in the same location in two separate cell lines and to have a similar level of sub-clonality (i.e., similar copy number).

### Permutation methodology

In order to determine the significance of overlap between replication timing variants and various other genomic features, we permuted the replication timing variant locations, and reciprocally permuted the locations of the genomic feature of interest. Significance was determined by comparing the overlap between variants and the tested feature of interest to 1000 permutations.

For generating permuted variant regions, we required the following:

1. Each permutation consisted of a number of permuted windows equal to the number of variants, and each permuted window was the same size as the variant from which it was derived.
2. Replication timing in the middle of the permuted windows was required to be within +/− 0.2 SD of the variant from which it was derived.
3. Permuted regions could not overlap variants or each other. Permuted regions were ordered by the p-value of the corresponding variant, so the most significant variant was permuted first, and later regions within this same permutation could not overlap the regions that were determined prior.
4. Permuted regions could not overlap gaps in the reference genome.
5. Regions had to have data in at least 50% of the replication timing bins.

For comparison to mouse UR regions and reactivated-X replication timing variants, we also performed reciprocal permutations, in which replication timing and size were retained relative to the those regions, rather than to the haploid replication timing variants. For quiescence-controlled permutations, we also required that each permutation was roughly the same percentage (+/− 5%) quiescent (defined by the 18-state ChromHMM model) as the matched replication timing variant.

In order to determine overlap, we considered the span of each replication timing variant. For example, if 40% of a replication timing variant coincided with a region of interest, the contribution of this region to the total overlap would be 0.4. Summed over all the variants, this meant that the total possible overlap for n variants ranged from 0 to n. Doing so normalized the contribution of each variant equally, regardless of size. Overlap between variants and a given dataset were then compared to the distribution of 1000 permutations for both variant region permutations and reciprocal permutations, and a two-tailed p-value was calculated from the z-score of the real overlap. For genes and histone marks, we considered these as discrete calls rather than regions, therefore overlap distributions considered the number of occurrences within variants rather than the variant span. For ChromHMM states, overlap was not normalized by variant size, and was instead calculated as the total number of base pairs in a given ChromHMM state; this was done to avoid the possibility that the contribution of important but physically small chromatin regions were deflated in large variants.

### Gene ontology

We used the enrichment analysis tool from the Gene Ontology Consortium (The Gene Ontology, 2019) to examine the genes in each variant class for biological process, molecular function, and cellular component enrichment. A Fischer exact test was used to calculate a false discovery rate, which we report for all ontologies.

### Mouse UR

Mouse underrepresented regions in TGCs were obtained from (Hannibal et al., 2014) and lifted-over from the mm9 mouse reference genome to the human genome reference hg19 using the UCSC liftover tool. There were 47 regions found in all mouse placenta cells. We took both the minimal region found across all six TGC samples, and the maximal regions which were found in the TGC sample taken at the latest stage of development. In both cases, 45 of the 47 regions were successfully lifted-over.

## Supplementary tables

**Table S1:**
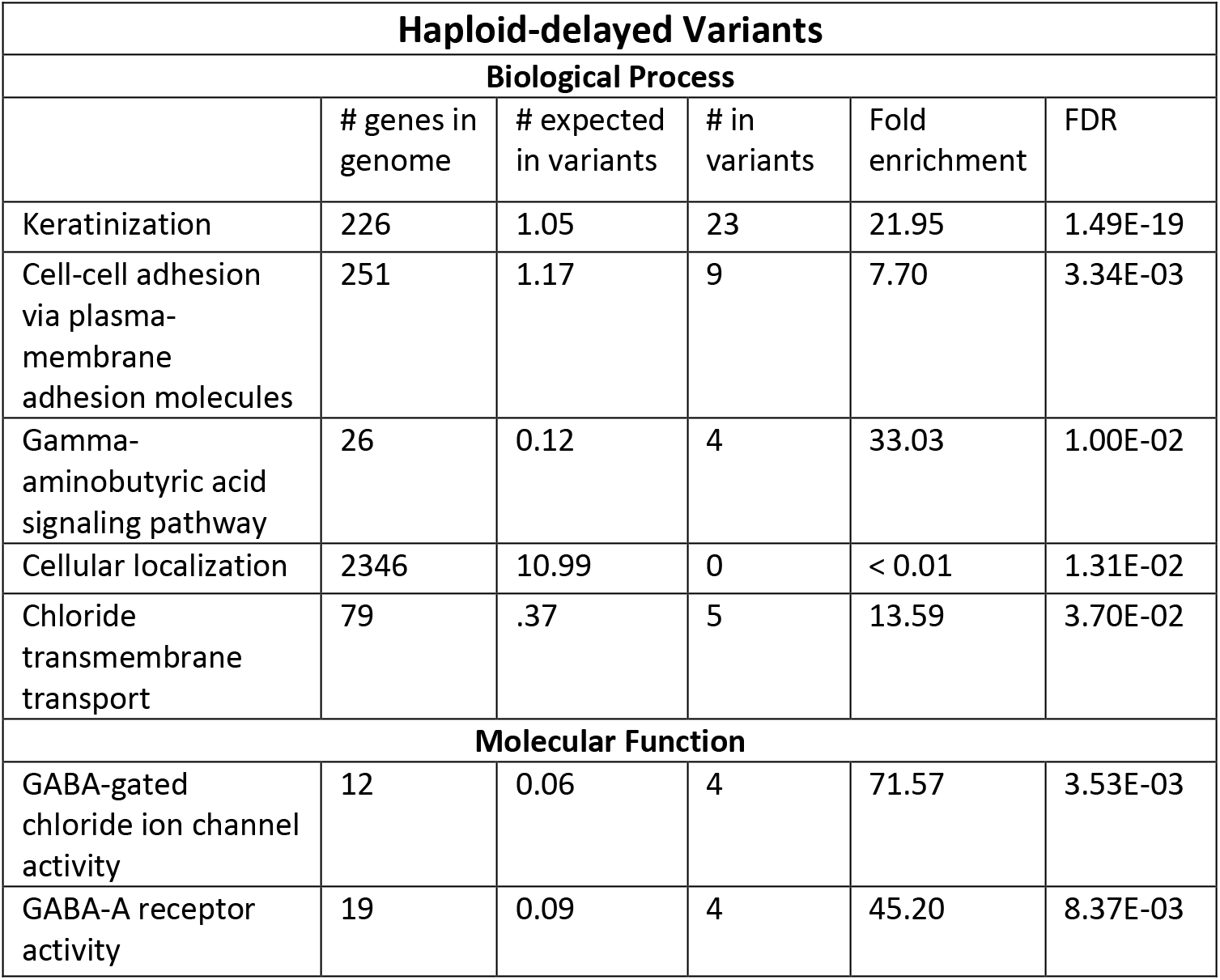
Results of GO analysis for genes found in variant regions. Nested groups were collapsed into only the most significant classification, such that when one biological process was a subset of another, and both were significant, only the most significant process is listed. **Haploid-delayed variants**: 97 of 180 genes were found in the GO database. **Haploid-advanced variants.** 41 of 168 genes were found in the GO database. **Downregulated genes found in haploid-advanced variants.** 11 of 13 genes were found in the GO database.

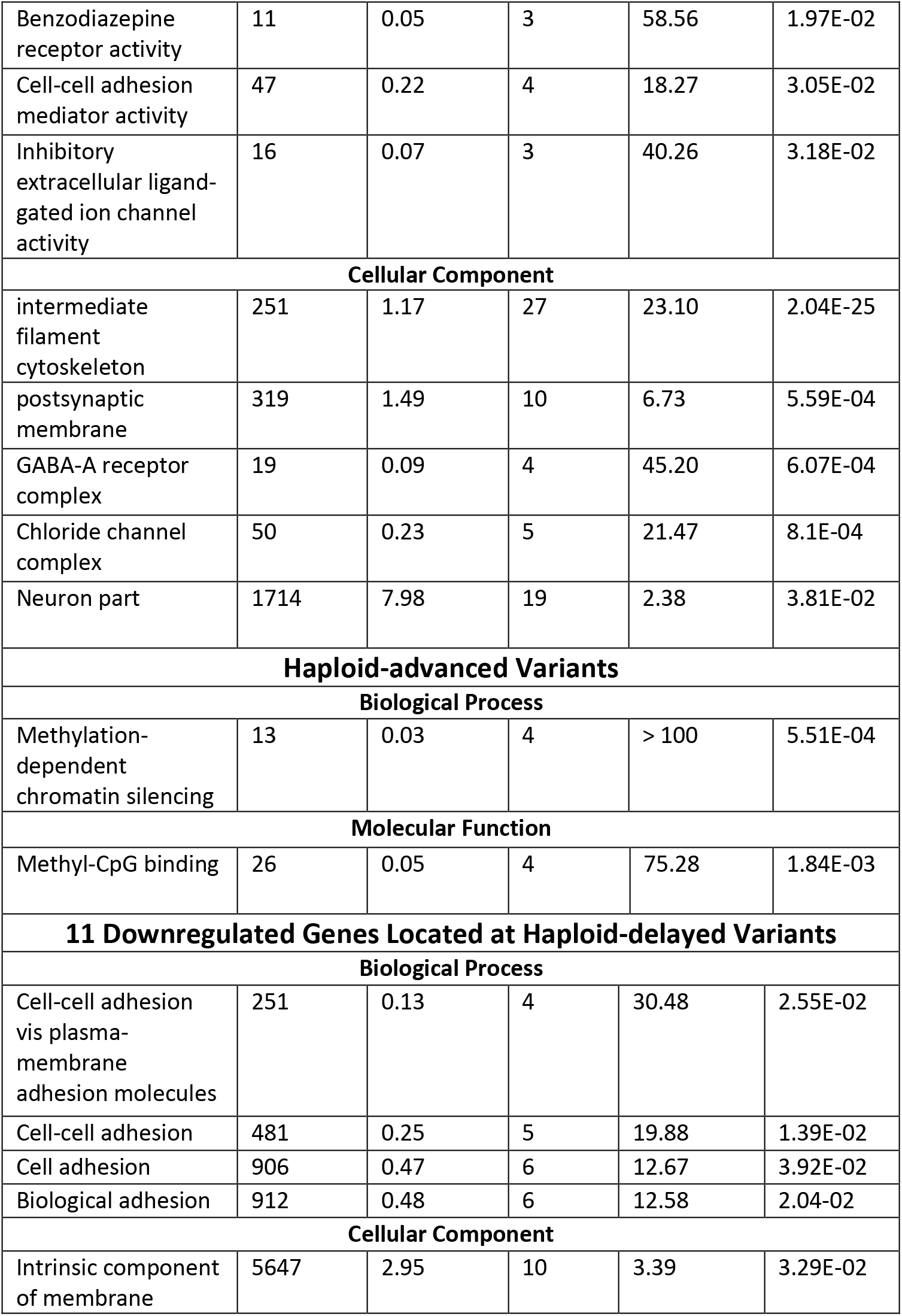

**Table S2.** Haploid-delayed variants. Shown are each of the 23 haploid-delayed variants with their variant number, genomic position in hg19 coordinates, and p-values in the initial and the two validation experiments including the recently diploidized cell lines. Significant p-values (p < 2.38 × 10^−3^; Bonferroni correction for 21 tests) are shown in bold. P-values for experiment 2 were calculated using a t-test comparing a given cell line to d-pES12 at the haploid-delayed variant region. P-values for experiment 3 were based on ANOVA tests of both haploid cell lines in comparison to diploid ESCs. Start and end position for X chromosome variants are esitmates based on the points at which haploid replication timing profiles become later than diploid. Because of the doasage imbalance between haploid and diploid cells on the X chromosome, these reigons are not amenable to the ANOVA test.

| Variant number | Chr | Start | End | Exp1 p (ANOVA) | h-pES10-exp2 p (t-test) | h-pES12-exp2 p (t-test) | RD-pES10-exp2 p (t-test) | RD-pES12-exp2 p (t-test) | haploid-exp3 p (ANOVA) | RD-exp3 p-value (ANOVA) |
| --- | --- | --- | --- | --- | --- | --- | --- | --- | --- | --- |
| 1 | 1 | 105036062 | 107043787 | **5.57E-50** | **4.96E-08** | **2.69E-20** | **4.22E-10** | **2.30E-25** | **5.63E-24** | **1.53E-09** |
| 2 | 1 | 188051349 | 192033027 | **3.43E-188** | **4.79E-11** | **3.29E-39** | **7.26E-22** | **8.02E-66** | **5.37E-71** | **3.35E-31** |
| 3 | 1 | 192862835 | 193254343 | **7.90E-06** | 0.0586 | 0.0063 | 0.668 | 0.949 | 0.0356 | 0.123 |
| 4 | 1 | 231571593 | 232457523 | **2.81E-10** | 0.203 | 0.393 | 0.662 | **6.06E-07** | 0.473 | 0.0896 |
| 5 | 2 | 13574856 | 14708037 | **3.11E-30** | **9.02E-06** | **1.73E-13** | 0.00909 | **3.77E-15** | **5.47E-16** | **8.38E-07** |
| 6 | 3 | 294211 | 2679244 | **6.71E-92** | **2.44E-18** | **1.49E-34** | **3.65E-11** | **2.73E-69** | **1.64E-70** | **5.39E-77** |
| 7 | 4 | 43409355 | 48656695 | **2.22E-76** | **5.52E-27** | **1.78E-63** | **1.54E-36** | **2.95E-77** | **3.28E-55** | **1.26E-78** |
| 8 | 4 | 59796831 | 65081128 | **9.65E-89** | **9.40E-39** | **1.54E-62** | **2.24E-39** | **3.63E-138** | **1.48E-55** | **2.18E-66** |
| 9 | 5 | 22493289 | 30383749 | **4.48E-184** | **3.81E-12** | **1.28E-84** | **1.71E-22** | **2.28E-165** | **9.57E-122** | **3.06E-188** |
| 10 | 6 | 62248453 | 68299337 | **1.21E-113** | **5.87E-22** | **2.14E-59** | **6.11E-36** | **3.04E-143** | **3.79E-150** | **2.28E-175** |
| 11 | 8 | 3393305 | 5944692 | **8.62E-60** | **5.05E-24** | **4.11E-31** | **1.99E-14** | **2.47E-64** | **1.23E-46** | **1.38E-40** |
| 12 | 8 | 13728440 | 15002424 | **8.67E-12** | **6.69E-15** | **8.97E-17** | **2.43E-10** | **2.29E-23** | **1.14E-24** | **4.88E-42** |
| 13 | 8 | 48348584 | 49252651 | **2.27E-10** | 0.134 | 0.0971 | 0.0265 | 0.83 | 0.205 | **5.48E-11** |
| 14 | 9 | 104921528 | 106545385 | **2.04E-51** | 0.0055 | **5.12E-14** | 0.00202 | **1.87E-27** | **2.13E-15** | **6.32E-18** |
| 15 | 10 | 108818724 | 111007775 | **2.06E-41** | **1.96E-09** | **1.77E-25** | **6.37E-12** | **6.19E-36** | **3.11E-22** | **5.07E-41** |
| 16 | 11 | 37239978 | 41263106 | **4.45E-68** | **3.62E-09** | **4.52E-23** | **5.90E-12** | **6.29E-81** | **1.62E-66** | **2.91E-54** |
| 17 | 12 | 127071746 | 127843422 | **1.61E-12** | **1.19E-07** | **9.03E-11** | **2.59E-08** | **9.39E-16** | **3.41E-08** | **1.70E-14** |
| 18 | 18 | 68680542 | 69909012 | **1.14E-41** | **2.04E-06** | **5.38E-18** | **4.04E-06** | **2.61E-19** | **8.48E-21** | **9.89E-13** |
| 19 | 20 | 53569880 | 54262237 | **6.22E-13** | 0.528 | **3.25E-08** | 0.813 | **3.08E-06** | **1.36E-17** | **5.10E-12** |
| 20 | 21 | 15376752 | 26722800 | **2.49E-302** | **1.91E-51** | **1.02E-99** | **7.81E-61** | **4.92E-249** | **< 2.5e-302** | **6.84E-193** |
| 21 | 21 | 31279591 | 32084063 | **3.47E-19** | 0.191 | **3.93E-08** | 0.498 | **1.62E-05** | **1.01E-23** | **6.08E-26** |
| 22 | X | 80992131 | 82965996 | N/A | N/A | N/A | N/A | N/A | N/A | N/A |
| 23 | X | 141643047 | 146459002 | N/A | N/A | N/A | N/A | N/A | N/A | N/A |

## References

Attia, M., Rachez, C., De Pauw, A., Avner, P., and Rogner, U.C. (2007). Nap1l2 promotes histone acetylation activity during neuronal differentiation. Mol Cell Biol 27, 6093–6102.

Baker, B.A., Frickey, L., Yu, I.T., Hawkins, E.P., Cushing, B., and Perlman, E.J. (1998). DNA content of ovarian immature teratomas and malignant germ cell tumors. Gynecol Oncol 71, 14–18.

Barlow, J.H., Faryabi, R.B., Callen, E., Wong, N., Malhowski, A., Chen, H.T., Gutierrez-Cruz, G., Sun, H.W., McKinnon, P., Wright, G., et al. (2013). Identification of early replicating fragile sites that contribute to genome instability. Cell 152, 620–632.

Becker, K.A., Ghule, P.N., Therrien, J.A., Lian, J.B., Stein, J.L., van Wijnen, A.J., and Stein, G.S. (2006). Self-renewal of human embryonic stem cells is supported by a shortened G1 cell cycle phase. J Cell Physiol 209, 883–893.

Bhowmick, R., Minocherhomji, S., and Hickson, I.D. (2016). RAD52 Facilitates Mitotic DNA Synthesis Following Replication Stress. Mol Cell 64, 1117–1126.

Bielski, C.M., Zehir, A., Penson, A.V., Donoghue, M.T.A., Chatila, W., Armenia, J., Chang, M.T., Schram, A.M., Jonsson, P., Bandlamudi, C., et al. (2018). Genome doubling shapes the evolution and prognosis of advanced cancers. Nat Genet 50, 1189–1195.

Bignell, G.R., Greenman, C.D., Davies, H., Butler, A.P., Edkins, S., Andrews, J.M., Buck, G., Chen, L., Beare, D., Latimer, C., et al. (2010). Signatures of mutation and selection in the cancer genome. Nature 463, 893–898.

Bruck, T., Yanuka, O., and Benvenisty, N. (2013). Human pluripotent stem cells with distinct X inactivation status show molecular and cellular differences controlled by the X-Linked ELK-1 gene. Cell Rep 4, 262–270.

Chang, B.H., Smith, L., Huang, J., and Thayer, M. (2007). Chromosomes with delayed replication timing lead to checkpoint activation, delayed recruitment of Aurora B and chromosome instability. Oncogene 26, 1852–1861.

Corbel, C., Diabangouaya, P., Gendrel, A.V., Chow, J.C., and Heard, E. (2013). Unusual chromatin status and organization of the inactive X chromosome in murine trophoblast giant cells. Development 140, 861–872.

Dekker, J., Belmont, A.S., Guttman, M., Leshyk, V.O., Lis, J.T., Lomvardas, S., Mirny, L.A., O’Shea, C.C., Park, P.J., Ren, B., et al. (2017). The 4D Nucleome Project. Nature 549, 219–226.

Ding, Q., Edwards, M.M., Hulke, M.L., Bracci, A.N., Hu, Y., Tong, Y., Zhu, X., Hsiao, J., Charvet, C.J., Ghosh, S., et al. (2020). The Genetic Architecture of DNA Replication Timing in Human Pluripotent Stem Cells. bioRxiv.

Dixon, J.R., Selvaraj, S., Yue, F., Kim, A., Li, Y., Shen, Y., Hu, M., Liu, J.S., and Ren, B. (2012). Topological Domains in Mammalian Genomes Identified by Analysis of Chromatin Interactions. Nature 485, 376–380.

Enoch, T., Carr, A.M., and Nurse, P. (1992). Fission yeast genes involved in coupling mitosis to completion of DNA replication. Genes Dev 6, 2035–2046.

Eykelenboom, J.K., Harte, E.C., Canavan, L., Pastor-Peidro, A., Calvo-Asensio, I., Llorens-Agost, M., and Lowndes, N.F. (2013). ATR activates the S-M checkpoint during unperturbed growth to ensure sufficient replication prior to mitotic onset. Cell Rep 5, 1095–1107.

Fan, J.B., Surti, U., Taillon-Miller, P., Hsie, L., Kennedy, G.C., Hoffner, L., Ryder, T., Mutch, D.G., and Kwok, P.Y. (2002). Paternal origins of complete hydatidiform moles proven by whole genome single-nucleotide polymorphism haplotyping. Genomics 79, 58–62.

Francisco, F.O., and Lemos, B. (2014). How do y-chromosomes modulate genome-wide epigenetic States: genome folding, chromatin sinks, and gene expression. J Genomics 2, 94–103.

Fungtammasan, A., Walsh, E., Chiaromonte, F., Eckert, K.A., and Makova, K.D. (2012). A genome-wide analysis of common fragile sites: what features determine chromosomal instability in the human genome? Genome Res 22, 993–1005.

Guo, A., Huang, S., Yu, J., Wang, H., Li, H., Pei, G., and Shen, L. (2017). Single-Cell Dynamic Analysis of Mitosis in Haploid Embryonic Stem Cells Shows the Prolonged Metaphase and Its Association with Self-diploidization. Stem Cell Reports 8, 1124–1134.

Hamline, M.Y., Corcoran, C.M., Wamstad, J.A., Miletich, I., Feng, J., Lohr, J.L., Hemberger, M., Sharpe, P.T., Gearhart, M.D., and Bardwell, V.J. (2020). OFCD syndrome and extraembryonic defects are revealed by conditional mutation of the Polycomb-group repressive complex 1.1 (PRC1.1) gene BCOR. Developmental biology 468, 110–132.

Handsaker, R.E., Van Doren, V., Berman, J.R., Genovese, G., Kashin, S., Boettger, L.M., and McCarroll, S.A. (2015). Large multiallelic copy number variations in humans. Nat Genet 47, 296–303.

Hannibal, R.L., and Baker, J.C. (2016). Selective Amplification of the Genome Surrounding Key Placental Genes in Trophoblast Giant Cells. Curr Biol 26, 230–236.

Hannibal, R.L., Chuong, E.B., Rivera-Mulia, J.C., Gilbert, D.M., Valouev, A., and Baker, J.C. (2014). Copy number variation is a fundamental aspect of the placental genome. PLoS Genet 10, e1004290.

Harrigan, J.A., Belotserkovskaya, R., Coates, J., Dimitrova, D.S., Polo, S.E., Bradshaw, C.R., Fraser, P., and Jackson, S.P. (2011). Replication stress induces 53BP1-containing OPT domains in G1 cells. J Cell Biol 193, 97–108.

Heskett, M.B., Sanborn, J.Z., Boniface, C., Goode, B., Chapman, J., Garg, K., Rabban, J.T., Zaloudek, C., Benz, S.C., Spellman, P.T., et al. (2020). Multiregion exome sequencing of ovarian immature teratomas reveals 2N near-diploid genomes, paucity of somatic mutations, and extensive allelic imbalances shared across mature, immature, and disseminated components. Mod Pathol 33, 1193–1206.

Hoffman, M.M., Ernst, J., Wilder, S.P., Kundaje, A., Harris, R.S., Libbrecht, M., Giardine, B., Ellenbogen, P.M., Bilmes, J.A., Birney, E., et al. (2013). Integrative annotation of chromatin elements from ENCODE data. Nucleic Acids Res 41, 827–841.

Huang, C., Cheng, J., Bawa-Khalfe, T., Yao, X., Chin, Y.E., and Yeh, E.T.H. (2016). SUMOylated ORC2 Recruits a Histone Demethylase to Regulate Centromeric Histone Modification and Genomic Stability. Cell Rep 15, 147–157.

Hulke, M.L., Massey, D.J., and Koren, A. (2020). Genomic methods for measuring DNA replication dynamics. Chromosome Res 28, 49–67.

Koren, A., Handsaker, R.E., Kamitaki, N., Karlic, R., Ghosh, S., Polak, P., Eggan, K., and McCarroll, S.A. (2014). Genetic variation in human DNA replication timing. Cell 159, 1015–1026.

Koren, A., and McCarroll, S.A. (2014). Random replication of the inactive X chromosome. Genome Res 24, 64–69.

Leeb, M., Dietmann, S., Paramor, M., Niwa, H., and Smith, A. (2014). Genetic exploration of the exit from self-renewal using haploid embryonic stem cells. Cell Stem Cell 14, 385–393.

Leeb, M., Walker, R., Mansfield, B., Nichols, J., Smith, A., and Wutz, A. (2012). Germline potential of parthenogenetic haploid mouse embryonic stem cells. Development 139, 3301–3305.

Leng, L., Ouyang, Q., Kong, X., Gong, F., Lu, C., Zhao, L., Shi, Y., Cheng, D., Hu, L., Lu, G., et al. (2017). Self-diploidization of human haploid parthenogenetic embryos through the Rho pathway regulates endomitosis and failed cytokinesis. Sci Rep 7, 4242.

Li, H., and Durbin, R. (2010). Fast and accurate long-read alignment with Burrows-Wheeler transform. Bioinformatics 26, 589–595.

Li, H., Guo, A., Xie, Z., Tu, W., Yu, J., Wang, H., Zhao, J., Zhong, C., Kang, J., Li, J., et al. (2017). Stabilization of mouse haploid embryonic stem cells with combined kinase and signal modulation. Sci Rep 7, 13222.

Li, Y., and Shuai, L. (2017). A versatile genetic tool: haploid cells (Stem Cell Research & Therapy).

Linder, D., McCaw, B.K., and Hecht, F. (1975). Parthenogenic origin of benign ovarian teratomas. N Engl J Med 292, 63–66.

Lukas, C., Savic, V., Bekker-Jensen, S., Doil, C., Neumann, B., Pedersen, R.S., Grofte, M., Chan, K.L., Hickson, I.D., Bartek, J., et al. (2011). 53BP1 nuclear bodies form around DNA lesions generated by mitotic transmission of chromosomes under replication stress. Nat Cell Biol 13, 243–253.

Makunin, I.V., Volkova, E.I., Belyaeva, E.S., Nabirochkina, E.N., Pirrotta, V., and Zhimulev, I.F. (2002). The Drosophila suppressor of underreplication protein binds to late-replicating regions of polytene chromosomes. Genetics 160, 1023–1034.

Minocherhomji, S., Ying, S., Bjerregaard, V.A., Bursomanno, S., Aleliunaite, A., Wu, W., Mankouri, H.W., Shen, H., Liu, Y., and Hickson, I.D. (2015). Replication stress activates DNA repair synthesis in mitosis. Nature 528, 286–290.

Munden, A., Rong, Z., Sun, A., Gangula, R., Mallal, S., and Nordman, J.T. (2018). Rif1 inhibits replication fork progression and controls DNA copy number in Drosophila. Elife 7.

Nordman, J., Li, S., Eng, T., Macalpine, D., and Orr-Weaver, T.L. (2011). Developmental control of the DNA replication and transcription programs. Genome Res 21, 175–181.

Nordman, J.T., Kozhevnikova, E.N., Verrijzer, C.P., Pindyurin, A.V., Andreyeva, E.N., Shloma, V.V., Zhimulev, I.F., and Orr-Weaver, T.L. (2014). DNA copy-number control through inhibition of replication fork progression. Cell Rep 9, 841–849.

Patel, S., Bonora, G., Sahakyan, A., Kim, R., Chronis, C., Langerman, J., Fitz-Gibbon, S., Rubbi, L., Skelton, R.J., Ardehali, R., et al. (2017). Human embryonic stem cells do not change their X-inactivation status during differentiation. Cell Rep 18, 54–67.

Rantala, J.K., Edgren, H., Lehtinen, L., Wolf, M., Kleivi, K., Vollan, H.K., Aaltola, A.R., Laasola, P., Kilpinen, S., Saviranta, P., et al. (2010). Integrative functional genomics analysis of sustained polyploidy phenotypes in breast cancer cells identifies an oncogenic profile for GINS2. Neoplasia 12, 877–888.

Roadmap Epigenomics, C., Kundaje, A., Meuleman, W., Ernst, J., Bilenky, M., Yen, A., Heravi-Moussavi, A., Kheradpour, P., Zhang, Z., Wang, J., et al. (2015). Integrative analysis of 111 reference human epigenomes. Nature 518, 317–330.

Ryba, T., Hiratani, I., Lu, J., Itoh, M., Kulik, M., Zhang, J., Schulz, T.C., Robins, A.J., Dalton, S., and Gilbert, D.M. (2010). Evolutionarily conserved replication timing profiles predict long-range chromatin interactions and distinguish closely related cell types. Genome Res 20, 761–770.

Sagi, I., and Benvenisty, N. (2017). Haploidy in Humans: An Evolutionary and Developmental Perspective. Dev Cell 41, 581–589.

Sagi, I., Chia, G., Golan-Lev, T., Peretz, M., Weissbein, U., Sui, L., Sauer, M.V., Yanuka, O., Egli, D., and Benvenisty, N. (2016). Derivation and differentiation of haploid human embryonic stem cells. Nature 532, 107–111.

Sagi, I., De Pinho, J.C., Zuccaro, M.V., Atzmon, C., Golan-Lev, T., Yanuka, O., Prosser, R., Sadowy, A., Perez, G., Cabral, T., et al. (2019). Distinct Imprinting Signatures and Biased Differentiation of Human Androgenetic and Parthenogenetic Embryonic Stem Cells. Cell Stem Cell 25, 419–432 e419.

Savelyeva, L., and Brueckner, L.M. (2014). Molecular characterization of common fragile sites as a strategy to discover cancer susceptibility genes. Cell Mol Life Sci 71, 4561–4575.

Schiltz, R.L., Mizzen, C.A., Vassilev, A., Cook, R.G., Allis, C.D., and Nakatani, Y. (1999). Overlapping but distinct patterns of histone acetylation by the human coactivators p300 and PCAF within nucleosomal substrates. J Biol Chem 274, 1189–1192.

Schoenfelder, K.P., and Fox, D.T. (2015). The expanding implications of polyploidy. J Cell Biol 209, 485–491.

Stelzer, Y., Yanuka, O., and Benvenisty, N. (2011). Global analysis of parental imprinting in human parthenogenetic induced pluripotent stem cells. Nat Struct Mol Biol 18, 735–741.

Stevens, L., and Varnum, D. (1973). The development of teratomas from parthenogenetically activated ovarian mouse eggs (Developmental Biology: Academic Press Inc.), pp. 369–380.

Sudmant, P.H., Rausch, T., Gardner, E.J., Handsaker, R.E., Abyzov, A., Huddleston, J., Zhang, Y., Ye, K., Jun, G., Fritz, M.H., et al. (2015). An integrated map of structural variation in 2,504 human genomes. Nature 526, 75–81.

Takahashi, S., Lee, J., Kohda, T., Matsuzawa, A., Kawasumi, M., Kanai-Azuma, M., Kaneko-Ishino, T., and Ishino, F. (2014). Induction of the G2/M transition stabilizes haploid embryonic stem cells. Development 141, 3842–3847.

Tarkowski, A.K., Witkowska, A., and Nowicka, J. (1970). Experimental partheonogenesis in the mouse. Nature 226, 162–165.

The Gene Ontology, C. (2019). The Gene Ontology Resource: 20 years and still GOing strong. Nucleic Acids Res 47, D330–D338.

Uhlen, M., Fagerberg, L., Hallstrom, B.M., Lindskog, C., Oksvold, P., Mardinoglu, A., Sivertsson, A., Kampf, C., Sjostedt, E., Asplund, A., et al. (2015). Proteomics. Tissue-based map of the human proteome. Science (New York, NY) 347, 1260419.

van Kempen, L.C., Redpath, M., Elchebly, M., Klein, K.O., Papadakis, A.I., Wilmott, J.S., Scolyer, R.A., Edqvist, P.H., Ponten, F., Schadendorf, D., et al. (2016). The protein phosphatase 2A regulatory subunit PR70 is a gonosomal melanoma tumor suppressor gene. Sci Transl Med 8, 369ra177.

Wear, E.E., Song, J., Zynda, G.J., Mickelson-Young, L., LeBlanc, C., Lee, T.J., Deppong, D.O., Allen, G.C., Martienssen, R.A., Vaughn, M.W., et al. (2020). Comparing DNA replication programs reveals large timing shifts at centromeres of endocycling cells in maize roots. PLoS Genet 16, e1008623.

Yaffe, E., Farkash-Amar, S., Polten, A., Yakhini, Z., Tanay, A., and Simon, I. (2010). Comparative analysis of DNA replication timing reveals conserved large-scale chromosomal architecture. PLoS Genet 6, e1001011.

Yaguchi, K., Yamamoto, T., Matsui, R., Tsukada, Y., Shibanuma, A., Kamimura, K., Koda, T., and Uehara, R. (2018). Uncoordinated centrosome cycle underlies the instability of non-diploid somatic cells in mammals. J Cell Biol 217, 2463–2483.

Yilmaz, A., Braverman-Gross, C., Bialer-Tsypin, A., Peretz, M., and Benvenisty, N. (2020). Mapping Gene Circuits Essential for Germ Layer Differentiation via Loss-of-Function Screens in Haploid Human Embryonic Stem Cells. Cell Stem Cell 27, 679–691 e676.

Yilmaz, A., Peretz, M., Sagi, I., and Benvenisty, N. (2016). Haploid Human Embryonic Stem Cells: Half the Genome, Double the Value. Cell Stem Cell 19, 569–572.

Zhang, X.M., Wu, K., Zheng, Y., Zhao, H., Gao, J., Hou, Z., Zhang, M., Liao, J., Zhang, J., Gao, Y., et al. (2020). In vitro expansion of human sperm through nuclear transfer. Cell Research 30, 356–359.

Zhao, Y., and Garcia, B.A. (2015). Comprehensive Catalog of Currently Documented Histone Modifications. Cold Spring Harb Perspect Biol 7, a025064.

Zlotorynski, E., Rahat, A., Skaug, J., Ben-Porat, N., Ozeri, E., Hershberg, R., Levi, A., Scherer, S.W., Margalit, H., and Kerem, B. (2003). Molecular Basis for Expression of Common and Rare Fragile Sites. In Mol Cell Biol, pp. 7143–7151.

Zybina, E.V., and Zybina, T.G. (1996). Polytene Chromosomes in Mammalian Cells. In International Review of Cytology, K.W. Jeon, ed. (Academic Press), pp. 53–119.

